# Dual kinase inhibition provides early clues to restoration of neuroimmune homeostasis in an AAV-haSyn model of Parkinson’s disease

**DOI:** 10.64898/2026.11.06.764938

**Authors:** Wesley Chiang, Ivette Sandoval, Herman Li, Luis Daniel Bernal-Conde, Rene Squire, Madeline K. Jensen, Jack Schwartz, Erica Squire, Eric J. Wagner, Fredric Manfredsson, Harris A Gelbard

**Author notes:** Co-first authors (Wesley Chiang, Ivette Sandoval, Herman Li). Co-senior and Corresponding Authors.

## Abstract

**Background:** Previous studies of drugs targeting mixed linage kinase type 3 (MLK3) and/or leucine rich repeat kinase type 2 (LRRK2) in sporadic Parkinson’s disease (PD) have failed to meet clinical endpoints despite multiple lines of evidence implicating these kinases in pathogenesis. This has reduced enthusiasm for MLK3 and LRRK2 as druggable targets. Largely overlooked is the evidence that these targets remain valid but must be considered in combination to achieve a therapeutic outcome. This provides the rationale for this study.

**Methods:** We conducted a pilot study modeling early, pre-symptomatic PD with subsequent evaluation of therapeutic potential of a multi-kinase inhibitor (URMC-099) in alleviating disease pathology. Sprague Dawley rats received unilateral intranigral injections of low dose AAV-haSyn. After a two-week recovery period, all animals were administered either once-daily intraperitoneal (i.p.) injection of URMC-099 or its vehicle control until their terminal endpoint. Induction of early PD, and its alleviation with URMC-099, was evaluated by lateralized motor impairment using the cylinder test and immunohistochemical analysis of neuropathology in the substantia nigra (SN) and striatum. Finally, an *in vitro* microglial PD model exposed to aSyn pre-formed fibrils and treated with URMC-099 was used to further evaluate the mechanism underlying therapeutic activity.

**Results:** A unilateral model of early and progressive AAV-haSyn model for PD was used to demonstrate the therapeutic potential of a targeted multi-kinase inhibitor to attenuate PD-associated neuropathology and prevent the manifestation of lateral motor deficits. Immunohistochemical analysis revealed protection of neuronal cellular integrity along the nigral-striatal pathway with once-daily i.p. URMC-099. The mitigation of degenerative neuropathology was linked to targeted inhibition of MLK3 and LRRK2 and complemented by reduced levels of gliosis in astrocytes and microglia in the SN. Single-microglia segmentation analysis suggested URMC-099 treatment preserved dynamic effector state switching in microglia to enhance autophagic flux to reduce the overall burden of pathologic aSyn. Transcriptomic analysis of a complementary *in vitro* microglia model corroborated the retention of a homeostatic-like effector state with URMC-099 treatment that was unique to MLK3/LRRK2 dual inhibition.

**Conclusions:** Unilateral injection of low dose AAV-haSyn successfully resulted in a lateralized model of early, progressive PD. Dual inhibition of aberrant MLK3 and LRRK2 activity with once-daily i.p. URMC-099 therapy significantly mitigated the development of PD pathology and progression in an early manifest phenotype. Our results present a potential mechanism of therapeutic action dependent on the dual inhibition of MLK3 and LRRK2. This presents a novel therapeutic modality that may be beneficial to a broad population of etiologic PD patients.

**Graphical Abstract**

## Introduction

Parkinson’s disease (PD) has grown in notoriety as a disorder of not only genetic origins, but now also a consequence of a broader scope of idiopathic origins, such as following certain acute infections as well as exposure to environmental neurotoxicants (1–3). These disparate etiologies complicate our understanding of disease pathogenesis in PD, which now consists of a heterogeneous collection of neurodegenerative pathologies (4–7). This has also led to limited success in identifying early biomarkers (i.e., predating neurologic symptomology) that are strong correlates for a broad spectrum of patients (8–11). To date, only seed-amplification assays of alpha synuclein (αSyn) levels from spinal fluid samples have been proven to be a robust risk indicator and diagnostic aid for PD (12).

Not only has screening for etiological biomarkers been elusive, so has the development of effective therapies. Currently, levodopa-carbidopa therapy (i.e., Sinemet) is the mainstay of symptomatic treatment. However, this therapeutic route does not prevent disease progression, and high dosages of levodopa-carbidopa can potentiate both motor symptoms (e.g. dyskinesia) and non-motor symptoms such as loss of impulse control (13–16). To tackle this, there has been an increased investment in PD therapeutics designed to specifically inhibit proteins implicated in the regulation of key signaling modules in disease pathogenesis. These have included the 2007 PRECEPT trial, focused on regulating aberrant inflammation by targeted inhibition of mixed-lineage kinase 3 (MLK3) (17,18), and the current LUMA phase 2b trial that targets leucine-rich repeat kinase 2 (LRRK2) to repair dysfunctional autophagolysosomal activity (19–23).

Thus, we propose that targeted regulation of both MLK3 and LRRK2 may offer complementary therapeutic benefits and benefit a broader population of idiopathic PD patients. Based on this hypothesis, we developed URMC-099, a small-molecule multi-kinase inhibitor specific to LRRK2 and MLK3 with brain penetrance and drug-like properties to inhibit both target kinases (17,24,25). Previous use cases of URMC-099 alleviated proteasomal stress in preclinical rodent models for other neurodegenerative diseases and overall reduced neuroinflammation, both of which are key features of PD etiopathology (5,6,11,26,27). Our approach targets both LRRK2 and MLK3 as related substrates for the eventual activation of the JNK, p38-MAPK, and ERK 1/2 pathways (27–30). With the convergence of LRRK2/MLK3 pathways, this suggests that inhibition of one kinase may be insufficient to prevent pathologic loss of dopaminergic function in PD with increasing evidence for redundant mechanisms that may mediate MLK3 downstream signaling (31–33).

Here, we report the therapeutic effects of dual kinase inhibition of LRRK2 and MLK3 with URMC-099 in a pre-clinical rat model of early PD, with further mechanistic evaluation in an *in vitro* microglia culture model of PD-associated αSyn burden. A low-dose regimen for adeno-associated virus (AAV) overexpression of human αSyn (haSyn) was delivered to the SN of Sprague Dawley rats to recapitulate the peritoxic events that occur prior to overt neurodegeneration (34–36). Unilateral delivery of the vector allowed for hemispheric differences in neuropathology with eventual manifestation of lateral motor differences. Key hallmarks of early manifest PD – phosphorylation of αSyn and aggregate formation, nigral neurodegeneration, and gliosis – were greatly attenuated in rats given once-daily URMC-099 therapy. We link improved resilience against pathologic αSyn burden in neurons and microglia to targeted inhibition of MLK3 and LRRK2 by URMC-099. Single-cell segmentation of microglia morphologic analysis with αSyn burden and LRRK2 activity identified a preservation of dynamic effector state switching in microglia with URMC-099 treatment. Transcriptomic analysis of an *in vitro* model of synuclein burden in BV-2 microglia-like cells was used to evaluate the unique mechanistic consequences of dual kinase inhibition by URMC-099 that preserves effector state dynamicity to result in enhanced autophagic flux.

Our findings suggest a therapeutic benefit of dual inhibition of LRRK2 and MLK3 signaling in attenuating the progression of PD neuropathology.

## Results

### Daily administration of URMC-099 reduces the severity of dopaminergic degeneration

Shown in Figure 1A, unilateral injection of AAV-haSyn in Sprague Dawley (SD) rats was used to induce intranigral expression of a transgene for human αSyn. Progression of synuclein-mediated pathology was allowed to progress for 6 weeks to recapitulate an established pre-clinical model of early PD (35). Dual-kinase inhibition of LRRK2/MLK3 was initiated two weeks after stereotactic surgery, with rats randomly assigned to cohorts that received either daily intraperitoneal (i.p.) injections of 10 mg/kg URMC-099 or matched vehicle controls. Similar to our previous findings, rats not administered any therapeutic (i.e., the vehicle cohort) demonstrated early indications of lateralized motor deficits in a cylinder test for preferential forepaw usage (Figure 1B). The asymmetric paw usage (17% differential) was linked to the lateralized expression of AAV-associated haSyn in the lesion-bearing hemisphere (Figure 1C). This was rescued in rats that received once-daily URMC-099 (i.e., the therapeutic cohort), which demonstrated symmetric paw usage (p = 0.008).

**Figure 1.**
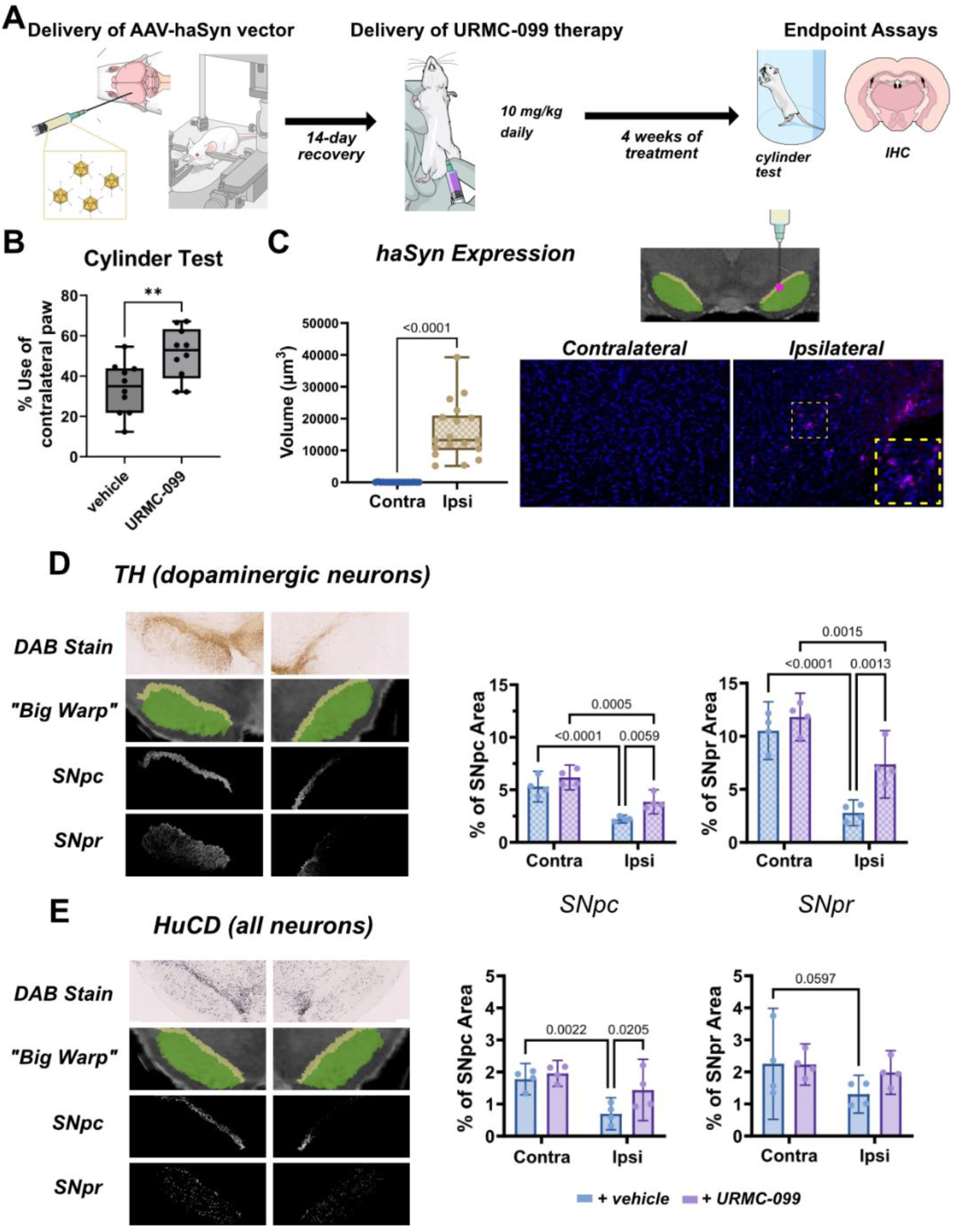
Induction of early PD pathology in unilateral AAV-haSyn paradigm and therapeutic attenuation by URMC-099. **(A)** Experimental outline of unilateral intranigral stereotactic injection of AAV-haSyn into Sprague Dawley rats (3-4 m.o.), dosing schedule of URMC-099 post-surgery, and endpoint (6-weeks post-injection) with associated assays. **(B)** Daily administration of 10 mg/kg URMC-099 demonstrates protection (p = 0.0081) against lateralized motor deficits in contralateral paw use as assessed via a cylinder test (n = 10 per therapeutic arm). **(C)** Diagram of localized delivery of AAV-haSyn on nigral-labeled Waxholm Space Atlas of SD brains. Quantification demonstrates significant (p < 0.0001) accumulation of αSyn in the ipsilateral SN with corresponding IHC panels of synuclein immunofluorescence that depict hemispheric expression (n = 18). **(D)** Schematic outline of sub-regional segmentation of nigral pars compacta and pars reticulata that was used to analyze changes in DAB immunoreactivity for dopaminergic neurons (TH+) and quantification of differences due to AAV-haSyn expression and URMC-099 treatment. **(E)** Segmentation and quantification for all neurons (HuC/D+) in indicative of overall neuropathology. In both populations, ipsilateral (i.e., lesion-bearing) hemispheres demonstrate decreases in neuronal coverage that are rescued with URMC-099 treatment. Plots in **B**-**E** reflect the mean and 95% confidence intervals. Statistical analyses for **B** and **C** were performed as two-tailed Mann-Whitney t-tests. Statistical analyses for **D** and **E** were performed as full effects two-way ANOVAs with Fisher’s LSD post-hoc corrections (n = 4 per category).

We then evaluated whether restoration of bilateral forepaw by URMC-099 treatment use was paralleled by a reduction in neurodegenerative pathology in the SN. High-resolution images were acquired from diaminobenzidine (DAB) stained midbrain sections immunolabeled for neuronal tyrosine hydroxylase (TH) and Hu antigens C/D (HuC/D) reactivity (Figures 1D and 1E). Annotated coronal reference sections from the Waxholm Space Atlas were used to align the images to the SN at matching coronal depths using the “Big Warp” plugin on ImageJ (37). DAB+ immunoreactivities in the pars compacta (SNpc, yellow) and pars reticulata (SNpr, green) were separately quantified to evaluate whether URMC-099 provided differential protection between the two nigral subregions, which reflect functionally distinct subregions of the basal ganglia (38–40). While the SNpr is primarily composed of GABAergic neurons, it is also a compartment densely innervated by the dendritic projections of the TH+ dopaminergic neurons that comprise the SNpc.

Because degeneration of dopaminergic neurons is a strong correlate of αSyn-associated neuropathology in PD (41), we first screened for differences the fractional area of the SN (i.e., % SN area) that is occupied by dopaminergic-specific TH+ immunoreactivity (Figure 1D). In the vehicle therapeutic arm, intranigral overexpression of hαSyn in the ipsilateral hemisphere significantly reduced the average % area that is TH+ in the SNpc by 3.1% (p < 0.0001) and the SNpr by 7.7% (p < 0.0001) compared the haSyn-free contralateral hemisphere. These mean differences amount to ∼60% and ∼80% less TH+ immunoreactivity in the ipsilateral SNpc and SNpr, respectively, when normalized to the mean % area that is TH+ in the contralateral subregions.

This vulnerability to αSyn of both nigral subregions was not recapitulated in our quantification of HuC/D immunoreactivity (Figure 1E), which captures overt neuronal loss as opposed to phenotypic loss of dopaminergic-specific TH immunoreactivity (42,43). A mean difference of 1.1% (p = 0.002) in HuC/D+ immunoreactive coverage of the SNpc was measured between the ipsilateral and contralateral hemispheres. This maps to a relative 61% reduction in HuC/D+ immunoreactivity in the ipsilateral SNpc, mirroring that of TH. In the SNpr, however, a mean difference of 0.9% (or relative reduction of 39%) between the two hemispheres in the percent coverage of subregion by HuC/D+ immunoreactivity was not found to be statistically significant (p = 0.06). The SNpr is predominantly populated by GABAergic neurons intermixed with a sparser population of dopaminergic projections. Thus, the loss of TH+ immunoreactivity across both subregions and SNpc-specific loss of HuC/D+ immunoreactivity confirms a distinct vulnerability of dopaminergic neurons to synuclein-mediated degeneration in our AAV-haSyn model.

Because rats given daily treatment with URMC-099 retained symmetrical usage of both forepaws, we then screened for whether this correlated with reduced neuropathology. Consequently, we observed significantly reduced dopaminergic degeneration in the ipsilateral SN. Compared to their vehicle cohort counterparts, rats given once-daily URMC-099 exhibited improved percent coverage of both the SNpc and SNpr by TH+ immunoreactivity by 1.6%, or a 1.5x increase, (p = 0.006) and 4.6%, or a 2.6x increase, (p = 0.001), respectively. Similar improvements were measured in the percent coverage of the SNpc by pan-neuronal HuC/D+ immunoreactivity (0.7% mean difference, or 2x increase, p = 0.021). No differences in the number of dopaminergic neurons, evaluated by enumeration of TH+ cell bodies, was observed (Supplemental Figure S1A). We also observed no differences in the average intensity per unit volume, a semi-quantitative proxy for protein expression levels, in neither TH+ nor HuC/D+ immunoreactive regions (Supplemental Figure S1B).

### URMC-099 diminishes phosphorylation burden on MLK3 and LRRK2

As diagrammed in Figure 2A, nigral neuroinflammation due to the accumulation of misfolded αSyn is expected to result in increased phosphorylation of MLK3 and LRRK2. Upregulation of the phosphorylated kinases (i.e., pMLK3 and pLRRK2) are correlated with activation of signaling cascades that promote progression of disease pathology (27,30,33,44,45). In previous rodent models of neuroinflammatory disease, we have demonstrated that MLK3 and LRRK2 are target substrates for URMC-099 with i.p. doses of 10 mg/kg resulting in reduction of pMLK3 and pLRRK2 levels in the brain (17,25,46). Thus, we evaluated changes in the overall phosphorylation burden on MLK3 (Figure 2B) and LRRK2 (Figure 2C) to confirm that the anti-inflammatory effect from URMC-099 treatment stems from the expected targeting of these two substrates.

**Figure 2.**
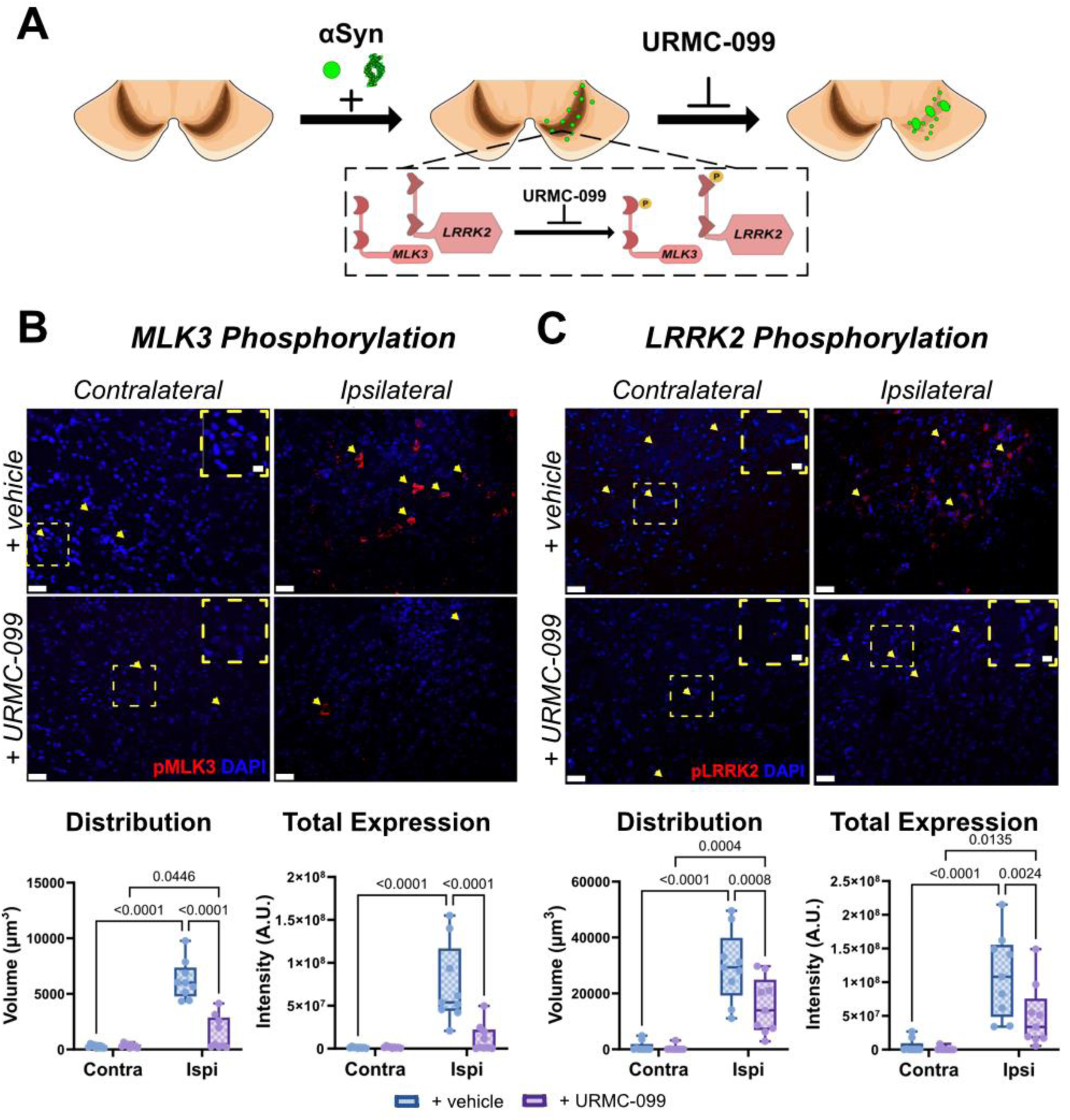
Overall burden of nigral pMLK3 and pLRRK2 reduced in rats given once-daily URMC-099. **(A)** Cartoon representing disease-associated upregulation in phosphorylation of MLK3 and LRRK2 following accumulation of αSyn. **(B)** Immunofluorescent staining of pMLK3 (red) in both hemispheres of AAV-haSyn SD rats given either vehicle of URMC-099 therapy. Yellow arrows serve to help identify regions of positive signal, with callout boxes to zoom in on smaller features. Scale bar is 30 μm. Graphs show quantification of changes in nigral levels of pMLK3 based on coverage (i.e. total volume) and protein content (i.e. total intensity). Treatment with URMC-099 results in ∼4-fold reduction in coverage of the SN by pMLK3 (p < 0.0001) and ∼50-fold reduction in total signal intensity (p < 0.0001) of pMLK3 in the ipsilateral SN. **(C)** Immunofluorescent staining of pLRRK2 (red) in both hemispheres of AAV-haSyn SD rats given either vehicle of URMC-099 therapy. Yellow arrows serve to help identify regions of positive signal, with callout boxes to zoom in on smaller features. Scale bar is 30 μm. Quantification of changes in nigral pLRRK2 based on same metrics for pMLK3. Treatment with URMC-099 results in 2-fold reduction in overall coverage of the SN by pLRRK2 (p = 0.0008) and ∼3-fold reduction in total signal intensity of pLRRK2 (p = 0.0024). All statistical tests were performed as two-way ANOVAs with Fisher’s LSD post-hoc corrections (n = 8 per treatment) and plotted as box plots for mean and interquartile range, with whiskers spanning to min and max values.

Expression patterns of pMLK3 in the SN were captured in the contralateral and ipsilateral hemispheres in both the vehicle and therapy cohorts (Figure 2B). In the contralateral SN, pMLK3+ signal was almost imperceivable (yellow arrows and dashed callout boxes), indicating the lack of an inflammatory environment and in agreement with the widespread TH+ and HuC/D+ immunoreactivity in the synuclein-free hemisphere (Figure 1C-E). Conversely, the ipsilateral SN exhibited significant increases of pMLK3+ features in both vehicle-treated (p < 0.0001) and URMC-099-treated (p = 0.045) rats. Widespread coverage of the SN by pMLK3 immunoreactivity (i.e., total volume) was observed in rats of vehicle therapeutic arm (average volume of 6000 μm^3^), while rats given once-daily URMC-099 were found to have a 4x reduction (mean difference = 4500 μm^3^, p < 0.0001) burden of total pMLK3 volume (average value of 1500 μm^3^). The total levels of pMLK3 in the SN (i.e., total intensity) also demonstrated a 7x reduction (mean difference = 6e7 a.u., p < 0.0001) between vehicle (average intensity of 7e7 a.u.) and therapy (average intensity 1e7 a.u.) cohorts.

Similarly, pLRRK2 immunoreactivity (Figure 2C), showed elevated phosphorylation localized to the ipsilateral nigra. Accordingly, rats given URMC-099 demonstrated a diminished burden in 2x total volume (mean difference = 1.5e4 μm^3^, p = 0.0008) and 3x total intensity (mean difference = 6e7 a.u., p = 0.002) of pLRRK2 compared to rats given i.p. vehicle. The reduced phosphorylation burden of both MLK3 and LRRK2 demonstrate effective target engagement by URMC-099 that corroborates the observed prevention of lateralized motor deficits and severity of neuropathology in rats of the therapeutic cohort.

### MLK3/LRRK2 dual inhibition reduces pathologic αSyn burden and alleviates gliosis

We then inquired whether dual inhibition of MLK3 and LRRK2 by URMC-099 would result in reduced levels of pathologic αSyn aggregates. Inhibition of MLK3 reduces phosphorylation of propyl-isomerase 1 (47), which is linked to its nuclear translocation and catalytic function relevant to the formation of nuclear αSyn inclusions in dopaminergic neurons (48). Correspondingly the ipsilateral SN in both vehicle and therapeutic cohorts exhibited heightened accumulation of haSyn within the soma (Supplemental Figure S5), reflective of cellular aSyn inclusions (49,50). Separately, LRRK2 dysfunction is linked to accumulation of neurotoxic protein aggregates due to poor regulation of autophagy-lysosomal flux (51–53).

Thus, we stained for phosphorylation of serine 129 on αSyn (pSer129) as an indicator for the presence of pathologic αSyn aggregates. In both the vehicle and therapeutic cohorts, the spread (i.e., total volume) of pSer129 in the SN were similar between the two groups (Supplemental Figure 2a) in both the contralateral (mean difference = 4.7e4 μm^3^, p = 0.11) and ipsilateral (mean difference = 1.5e4 μm^3^, p = 0.60) hemispheres. On average, the contralateral SN demonstrated a sparser (mean difference = 1.9e5 μm^3^, p < 0.0001) coverage by pSer129 (average volume = 1.7e5 μm^3^) that is linked to a homeostatic, physiological role for the phosphorylated state (54,55). The pervasive coverage of the SN by pSer129 (average volume = 3.6e5 μm^3^) in the AAV-haSyn-injected ipsilateral hemisphere reflects the activation of an aSyn-mediated pathology.

While the coverage of the SN by pSer129 demonstrated hemispheric differences that were independent of therapeutic cohort, the overall expression levels (i.e., total intensity) of pSer129 were found to have treatment-dependent differences in the ipsilateral SN (Figure 3A). As expected, the contralateral SN demonstrated similar pSer129 expression levels (mean difference = 0.6e8 a.u., p = 0.80) between the vehicle (average intensity = 3.8e8 a.u.) and therapeutic (average intensity = 4.4e8 a.u.) cohorts that were on average significantly lower (mean difference = 13.8e8 a.u., p < 0.0001) than the average pSer129 levels in the ipsilateral SN. However, within the ipsilateral SN, the once-daily URMC-099 therapeutic cohort demonstrated 30% lower pSer129 levels (mean difference = 6.4e8 a.u., p = 0.01) than that of the vehicle cohort (average intensity = 21.1e8 a.u.). This alleviation in the overall pSer129 burden in the injected hemisphere of the therapeutic cohort aligns with our expectations of URMC-099-mediated inhibition of aberrant MLK3 and LRRK2 activity linked to worsened disease pathology.

**Figure 3.**
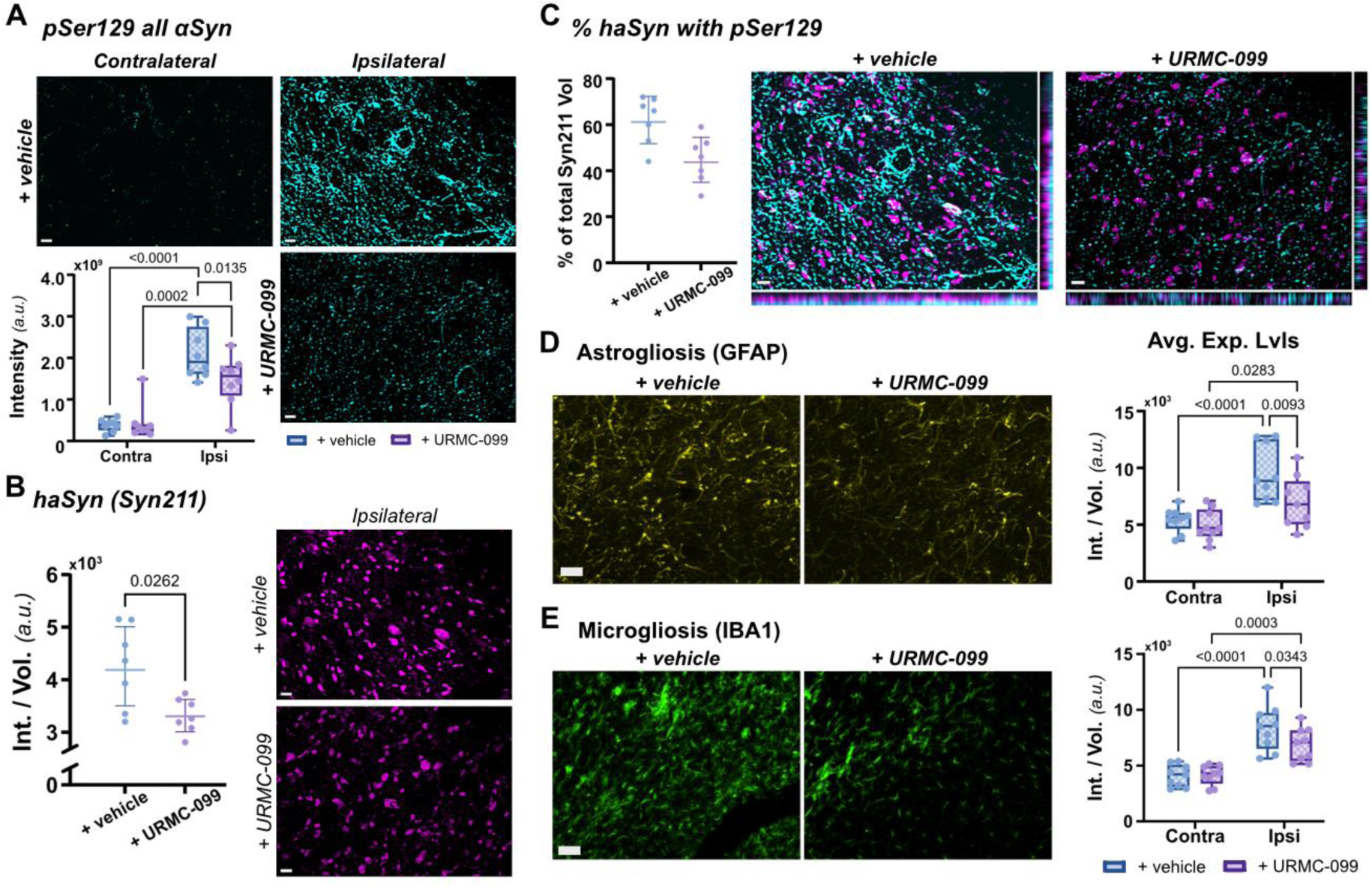
URMC-099 inhibition of MLK3 and LRRK2 reduces formation of pathologic αSyn aggregates and alleviates glial cell inflammation. **(A)** AAV-mediated expression of haSyn results in elevated αSyn phosphorylation (pSer129) in the ipsilateral hemisphere (∼5.5x greater than contralateral, p < 0.0001). Daily treatment with 10 mg/kg URMC-099 reduces total pSer129 levels by 30% (p = 0.0135) compared to daily vehicle treatment. Scale bar is 5 μm. **(B)** URMC-099 treatment also results in ∼20% lower average levels of haSyn (Syn211+) compared to rats in the vehicle therapeutic arm (p = 0.0262). Scale bar is 5 μm. **(C)** Colocalization analysis of pSer129+ with Syn211+ AAV-induced haSyn demonstrated a reduction in the fraction of total haSyn that is phosphorylated [(pSer129+ Syn211+) / Syn211+] from ∼60% to ∼45% (p = 0.007) with URMC-099 treatment. Scale bar is 5 μm. **(D)** IHC panels for GFAP immunofluorescence that are representative of changes in astroglial health. Scale bar is 30 μm. Significant (p < 0.0001) hemispheric differences in GFAP expression levels (60% greater in ipsilateral) as a result of hαSyn expression. URMC-099 treatment diminishes astrogliosis in ipsilateral SN by ∼25% (p = 0.0093). **(E)** IHC panels for IBA1 immunofluorescence representative of changes in microglial health. Scale bar is 30 μm. Significant (p < 0.0001) hemispheric differences in IBA1 expression levels (2.5-fold greater in ipsilateral) resultant from haSyn expression. URMC-099 treatment results in 10% reduction of astrogliosis. Statistical analyses for panels **B** and **C** were performed as two-tailed Mann-Whitney t-tests (n = 7 per treatment) and plotted as mean with 95% confidence intervals. Statistical analyses for A (n = 7 per treatment), **D** and **E** (n = 8 per treatment) applied full-effects two-way ANOVAs with Fisher’s LSD post-hoc corrections and plotted as box plots for mean and interquartile range, with whiskers spanning to min and max values.

The measured pSer129 burden captures phosphorylation of both endogenous rat αSyn and AAV-induced haSyn. Thus, we wanted to evaluate whether the haSyn species was actively participating in the formation of larger aggregates and inclusions (Supplemental Figure S5). Given the limited expression of haSyn in the non-injected hemisphere, we focused on evaluating the sequestration of Syn211+ haSyn into bright, localized aggregates (i.e., an increased average intensity per volume) in the ipsilateral hemisphere (Figure 3B). Compared to those given vehicle therapeutic, rats given once-daily URMC-099 therapy demonstrated a 20% reduction (difference = 1000 a.u., p = 0.03) in the average expression level of haSyn per Syn211+ feature (vehicle average = 4300 a.u., URMC-099 average = 3300 a.u.). This was complemented by a ∼2x sparser (difference = 2.8e5 μm^3^, p = 0.011) coverage of Syn211+ immunoreactivity throughout the ipsilateral SN (Supplemental Figure S2b) in the therapeutic cohort (average volume = 3.7e5 μm^3^) than the vehicle cohort (average volume = 6.5e5 μm^3^).

We also observed colocalization of pSer129 immunoreactivity with Syn211+ AAV-induced haSyn (Figure 3C), suggesting recognition of the human-derived αSyn by endogenous rat kinases as a phosphorylation substrate. The fraction of total haSyn that was phosphorylated (i.e., ratio of dual immunoreactive pSer129+ Syn211+ haSyn to all Syn211+ haSyn) was significantly lowered (p = 0.007) from ∼60% on average, in rats given vehicle therapeutic, to ∼45% on average, in rats given URMC-099. This further demonstrates that URMC-099-dependent regulation of LRRK2 signaling protects against the onset of pathologic signaling, such as aberrant αSyn phosphorylation leading to widespread aggregate formation.

Relatedly, aberrant activation of MLK3 and LRRK2 with the formation of pathologic αSyn inclusions not only affects neuronal health in the ipsilateral SN, but also has implications on the effector state of neighboring glial cells. To that end, we quantified levels of astrocyte and microglia activation using GFAP and IBA1 immunoreactivity, respectively (Figures 3D and 3E, Supplemental Figures S3 and S4). Regardless of therapeutic treatment, the average expression levels (i.e., intensity normalized by volume) of astrocytes (8e3 a.u.) and microglia (5e3 a.u.) were significantly elevated (p < 0.0001 for both) in the ipsilateral SN compared to cells in the contralateral SN (5e3 a.u. and 2e3 a.u., respectively). While GFAP and IBA1 expression levels were not restored to homeostatic levels in the ipsilateral SN of the therapeutic cohort, we still identified significant alleviation of both astrogliosis (mean difference = 2.5e3, p = 0.01) and microgliosis (mean difference = 1.0e3, p = 0.01) with once-daily URMC-099 treatment. Similar to the reduction in pathologic levels of pSer129+ αSyn aggregates in the ipsilateral SN by, we measured corresponding alleviation of gliosis would relieve the inflammatory stress imposed on astrocytes and microglia. We attribute the reduced severity of gliosis in the ipsilateral SN to be a combination of URMC-099 mediated reduction in pSer129+ αSyn aggregates and regulation of MLK3-dependent inflammatory signaling.

### URMC-099 treatment reinforces integrity of pre-synaptic terminal

The reduction in pathologic αSyn and related gliosis with URMC-099 treatment suggested the presence of a neuroprotective immune effector state in the SN. Given the high levels of αSyn in the synapse and the native role of αSyn in regulating synaptic vesicle activity (56,57), we sought to evaluate whether URMC-099-mediated neuroprotection was related to enhanced integrity of the pre-synaptic terminal by limiting the formation of pSer129+ αSyn aggregates.

Neurons in the SN were labeled for MAP2 and demonstrated differences in cellular integrity between the hemispheres (Figures 4A and 4B) that were consistent with our TH+ and HuC/D+ DAB analysis (Figures 1D and 1E). The regions outlined in Figure 4A were resampled to produce a zoomed in view (Figure 4C) to specifically evaluate cellular integrity along the neuronal projections. As shown in Figure 4B, neurons in the ipsilateral nigra of the vehicle cohort exhibited fragmented discontinuities along their processes, leading to ∼30% less MAP2+ coverage (mean difference = 6e4 μm^3^, p < 0.0001) of the SN compared to that in the contralateral SN (average volume = 26e4 μm^3^). Nigral neurons from rats of the therapeutic cohort exhibited healthier extended processes (mean difference = 7e4 μm^3^, p < 0.0001) that reflected more complete coverage of the ipsilateral SN by MAP2+ immunoreactivity (average volume = 27e4 μm^3^) than those from the vehicle cohort (average volume = 20e4 μm^3^).

**Figure 4.**
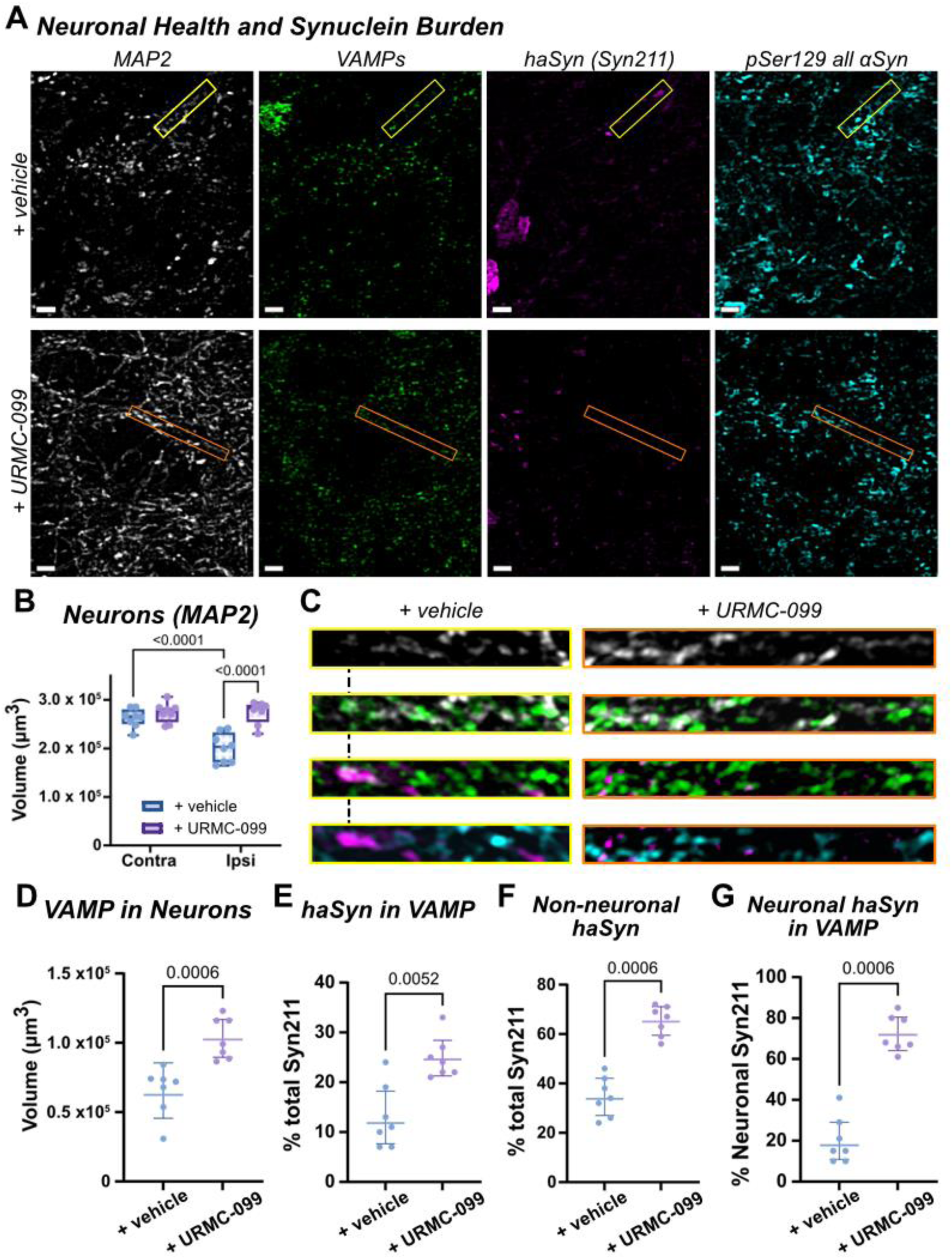
Effects of pathologic αSyn on neuronal cellular pathology. **(A)** Neurons in the ipsilateral SN were identified by MAP2 immunoreactivity and evaluated for the correlation between VAMP expression with pathologic αSyn burden along the neuronal processes. Scale bar = 5 μm. **(B)** Quantification of changes in coverage of SN by MAP2+ projections, demonstrating a ∼30% loss in the ipsilateral hemisphere of rats given vehicle therapy (p < 0.0001). This effect is significantly attenuated (p < 0.0001) in the ipsilateral SN of rats given URMC-099. **(C)** Zoomed in regions resampled that correspond to the outlined segments along a neuronal projection in **A**. Dashes between the zoomed in images of the +vehicle therapy set highlight the association of a large Syn211+ cluster with a loss of MAP2 intensity and lack of VAMP co-expression. **(D)** Quantification of total VAMP expression in MAP2+ neurons in the ipsilateral SN from the two therapeutic arms. URMC-099 treatment results in ∼1.8x greater distribution of VAMP along neurons (p = 0.0006). **(E)** Quantification of the fraction of all Syn211+ objects contained within neuronal VAMP features. Rats given URMC-099 demonstrate ∼2.2x more haSyn in VAMP (p = 0.0052). **(F)** Quantification of Syn211+ MAP2+ features identified non-neuronal haSyn immunoreactivity in the ipsilateral SN for both vehicle-treated (35% of all Syn211+ objects) and URMC-099 (∼70% of all Syn211+ objects), a 2x higher frequency of non-neuronal haSyn in rats given URMC-099 (p = 0.0006). **(G)** Correcting for Syn211+ VAMP+ colocalization based on MAP2+ Syn211+ objects demonstrates that URMC-099 treatment has a 3.5x enhancement of neuronal haSyn colocalization with VAMP (p = 0.0006). Statistical analyses for panels **D-G** were performed as unpaired two-tailed t-tests (n = 7 per treatment) and plotted as mean with 95% confidence intervals. Statistical analysis for **B** was performed as a full-effects two-way ANOVA with Fisher’s LSD post-hoc corrections (n = 7 per treatment) and plotted as box plots for mean and interquartile range, with whiskers spanning to min and max values.

This resulted in non-significant differences in neuronal health and integrity between the two hemispheres in the SN of rats from the therapy cohort. We also queried whether increased resilience of nigral dopaminergic neurons extended along the nigral-striatal pathway (Supplemental Figure 6). Specifically, we evaluated the ratio of total counts of Homer1+ post-synaptic densities between the two hemispheres as an indicator for changes in dopamine-regulated corticostriatal glutamate transmission reported in αSyn overexpression models (40,58,59). Treatment with URMC-099 significantly (p = 0.0072) improved the retention near similar levels striatal Homer1+ synapses between the hemispheres (average ratio = 96%) from the 76% ipsilateral to contralateral ratio in the striatum of vehicle-treated rats.

The health of pre-synaptic terminals along MAP2+ processes was then evaluated based on their αSyn pathology in relation to vesicle-associated membrane protein (VAMP) co-expression (Figures 4A and 4C). VAMPs are not only important to synaptic plasticity (56,60), but the VAMP2 isoform has also been identified as a chaperone of αSyn that regulates its condensation and aggregation potential (57,61). Thus, we screened for changes in VAMP expression in nigral neurons only in the ipsilateral hemisphere to screen for how these changes correlate to haSyn expression patterns. Nigral neurons in the ipsilateral hemisphere of the vehicle cohort exhibited ∼30% reduced coverage (difference = 2.7e4 μm^3^, p = 0.0006) of VAMP along their MAP2+ processes (average volume = 6.5e4 μm^3^) compared to the nigral neurons of the therapeutic cohort (average volume = 10.3e4 μm^3^). Zooming in on an individual MAP2+ neuronal process with haSyn (Syn211+) and pSer129+ aggregates (Figure 4C), we observed that these highly concentrated regions of pathologic αSyn to occupy VAMP deficient regions. In nigral neurons of the vehicle cohort, an overlap of only 13% was quantified between haSyn and VAMP (Figure 4E). The overlap between the two in MAP2+ processes improved ∼2x (difference = 12%, p = 0.005) in nigral neurons of the therapeutic cohort, and resulted in smaller, more punctal aggregates of Syn211+ and pSer129+ (Figures 4C and 4E).

However, the low relative fraction of total haSyn in the URMC-099 cohort (< 25% of all Syn211+ volume) led us to evaluate whether a sizable non-neuronal fraction of haSyn was present (Figure 4F, Supplemental Figure S7). A distinct (p = 0.0006) fraction of non-neuronal Syn211+ was quantified between the ipsilateral nigra of the two cohorts, reflecting ∼35% of all haSyn in the vehicle cohort and ∼70% in that of the URMC-099 therapeutic cohort. To account for colocalization between haSyn and VAMP that can only occur in neurons (i.e., haSyn+ VAMP+ MAP2+ features), we normalized volume of this population against the total volume of Syn211+ MAP2+ features (Figure 4G). Vehicle-treated rats still demonstrated a low colocalization of neuronal haSyn in VAMP (∼20%), but colocalization of neuronal haSyn with VAMP significantly (p = 0.0006) increased to ∼70% of all neuronal haSyn. These results suggest retention of homeostatic, co-regulatory activity between VAMP and αSyn that promotes physiological function rather than progression towards a neuropathological state.

### Alleviation in LRRK2 phosphorylation burden enhances microglial processing, but not engulfment, of αSyn

We then queried whether the non-neuronal fraction of haSyn in both treatment cohorts reflects microglia-mediated phagocytosis of stressed synaptic terminals, similar to that observed by amyloid-burdened synapses in Alzheimer’s Disease models (62,63). In our previous studies, we found that treatment with URMC-099 enhanced the capacity of autophagolysosomal processing in microglia to reduce the total amyloid burden in the hippocampus in a murine model of Alzheimer’s Disease (24). Thus, we tested whether a similar role for microglia-mediated degradation of pathologic αSyn could be enhanced by URMC-099 in our PD model.

Volumetric images of Tmem119+ microglia were segmented using the Volocity Quantification Suite to perform single cell analysis of LRRK2 and pLRRK2 expression levels (Figure 5A). This found a 24% increase (p < 0.0001) in the fraction of Tmem119+ microglia with elevated LRRK2 expression in the ipsilateral SN (54% of cells on average) compared to Tmem119+ microglia in the contralateral SN (30% of cells on average), indicative of upregulated activity in the AAV-haSyn-injected hemisphere (Figure 5B). We then quantified population-level statistics for changes in LRRK2 phosphorylation (pLRRK2) across segmented microglia as a measure of autophagic flux. While URMC-099 treatment did not reduce the number of Tmem119+ microglia with elevated LRRK2 expression (mean difference between therapy cohorts = 9%, p = 0.05), it did reduce the phosphorylation burden of microglial LRRK2 by 40% (p = 0.048) compared to vehicle-treated rats (Figure 5C). The lower pLRRK2:LRRK2 ratio suggests attenuation of LRRK2 activity, which was expected to promote autophagy in the microglia (51,52,64,65).

**Figure 5.**
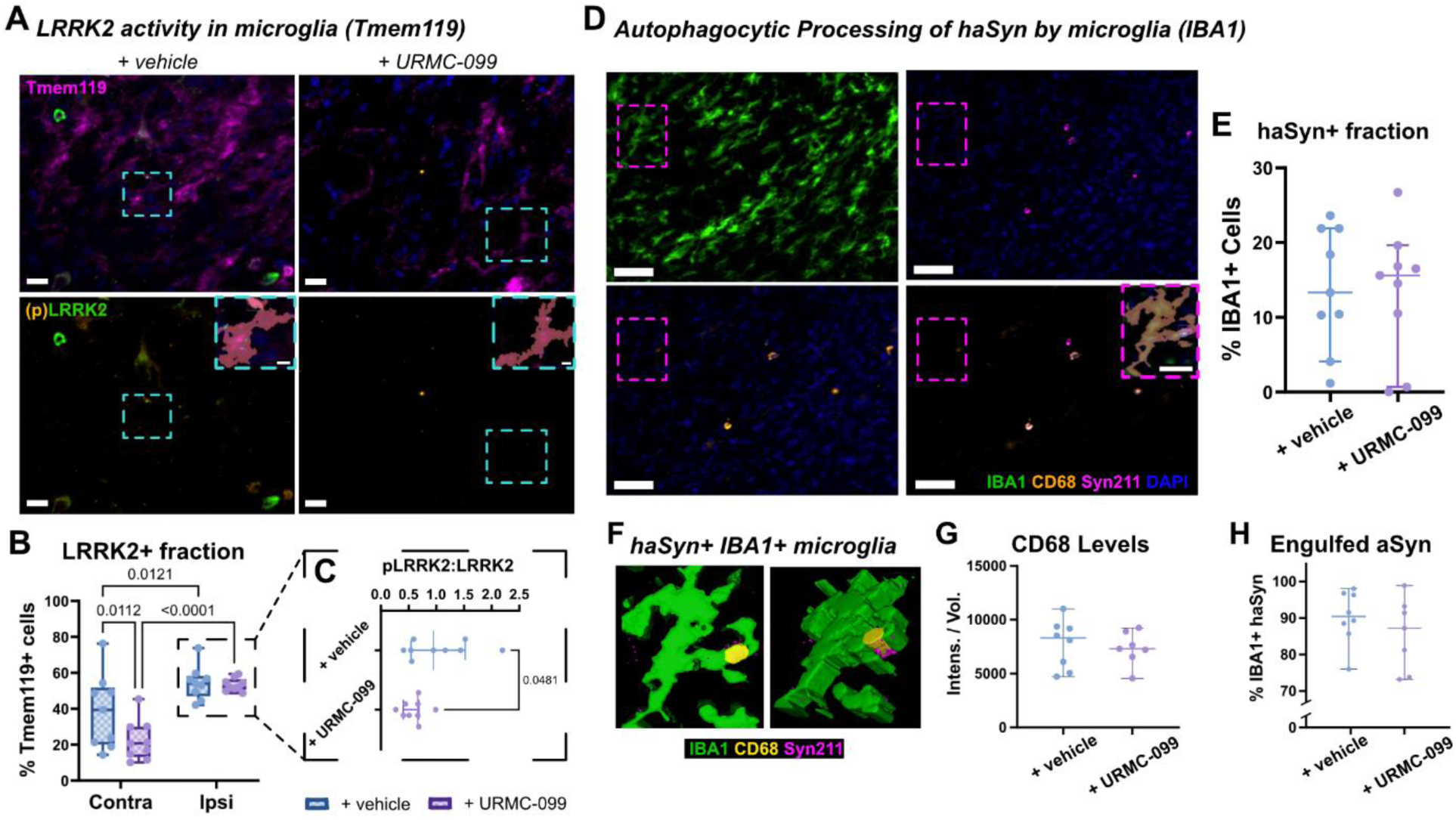
Autophagic processing of αSyn. **(A)** IHC staining and segmentation of Tmem119+ microglia in the ipsilateral SN of vehicle and URMC-099 treated rats. Example segmented microglia are enlarged as insets over the (p)LRRK2 channel, with a reddish-pink overlay depicting the Tmem119+ cell and with smaller objects highlighted to represent LRRK2 and pLRRK2. Scale bar is 15 μm. **(B)** Fraction of Tmem119+ microglia with elevated levels of LRRK2 expression. Regardless of therapy arm, microglia in the ipsilateral SN demonstrate heightened LRRK2 expression compared to the contralateral (vehicle: p = 0.0121; URMC-099: p < 0.0001). **(C)** In LRRK2+ Tmem119+ microglia in the ipsilateral SN, URMC-099 treatment reduces the activation of LRRK2, evaluated by pLRRK2 to LRRK2 ratio, by 50% compared to vehicle-treated rats (p = 0.048). **(D)** Representative IHC panel and segmentation of IBA1+ microglia in the ipsilateral SN to evaluate Syn211+ haSyn uptake and trafficking into CD68+ lysosomes. Scale bar 20 μm. Insert shows representative IBA1+ microglia with contained Syn211+ and CD68+ objects. Scale bar 10 μm. **(E)** Treatment with URMC-099 did not result in differences in the fraction of IBA1+ cells that contain haSyn. **(F)** Representative volumetric rendering of segmented IBA1+ microglia with Syn211+ haSyn contained with a CD68+ structure. No differences in **(G)** the average expression levels of CD68 nor **(H)** fraction of IBA1+ haSyn engulfed within CD68 structures. Statistical analyses for panels **C, E, G,** and **H** were performed as unpaired two-tailed t-tests (n = 8 per treatment) and plotted as mean with 95% confidence intervals. Statistical analysis for **B** was performed as full-effects two-way ANOVAs with Fisher’s LSD post-hoc corrections (n = 8 per treatment) and plotted as box plots for mean and interquartile range, with whiskers spanning to min and max values.

Using a similar segmentation protocol, we then evaluated the colocalization of Syn211+ features within CD68+ lysosomes in IBA1+ microglia as a measure of autophagic processing of synuclein (Figure 5D). Only IBA1+ microglia in the ipsilateral SN demonstrated association with Syn211+ signal (Supplemental Figure S8), in line with the hemispheric localization of haSyn expression (Figure 1C). Surprisingly, no differences were observed (mean difference = 0.5%, p = 0.91) in the fraction of IBA1+ microglia containing haSyn between therapy cohorts (Figure 5E). Further evaluation of only Syn211+ microglia (Figures 5F-H) also found no differences in CD68 expression levels (mean difference = 500, p = 0.61) and trafficking, or colocalization, of haSyn into CD68+ lysosomal structures (mean difference = 3%, p = 0.72).

### Retention of dynamicity in microglia to transition between effector states is linked to enhanced autophagic flux

While no differences were observed in the average frequency of microglia engulfment of αSyn, rats in the therapeutic cohort demonstrated a clear reduction in both total and pathologic αSyn burden (Figures 3A-C). Thus, we queried whether population of microglia containing αSyn in the vehicle cohort reflected a state of phagocytic exhaustion with concomitant reductions in autophagolysosomal flux (66,67). To do so, we classified segmented microglia (Figure 6A) based on the degree of elevation IBA1 expression levels and standard 3D morphologic parameters reflective of known “activated” effector states (Supplemental Figure S9) (68,69). Changes in distribution of the different classifications served as a measure for how well the microglia retained their ability to dynamically transition between effector states.

**Figure 6.**
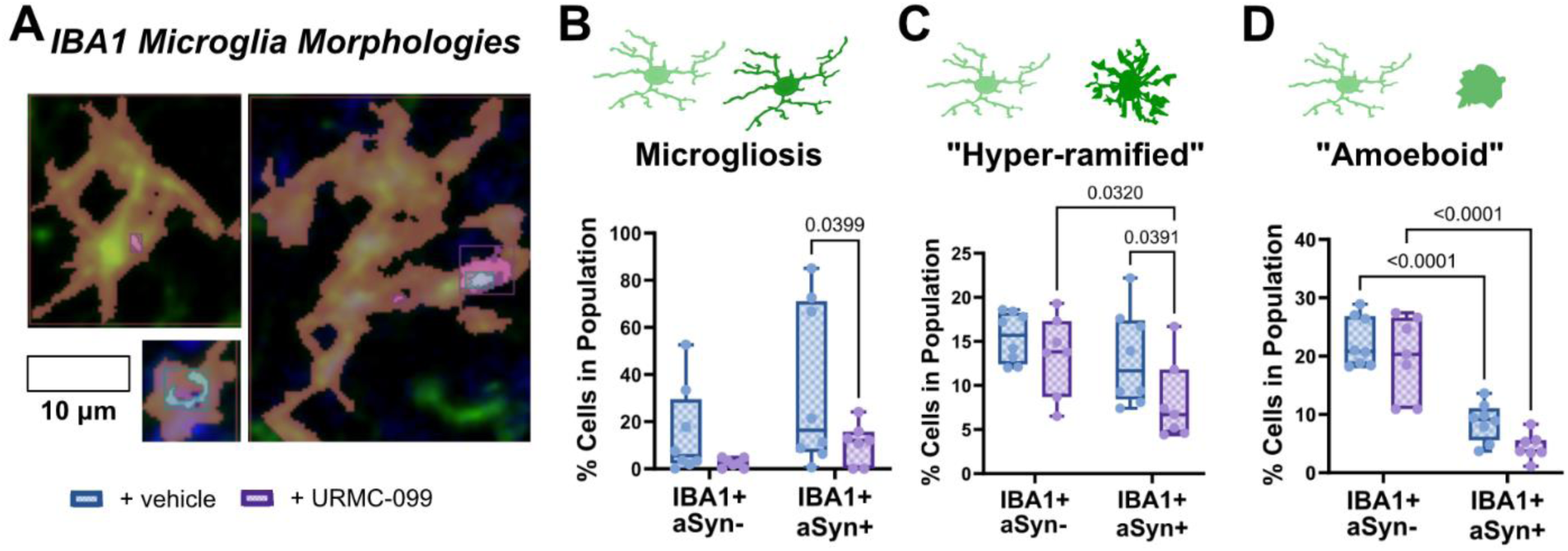
Effector state classification of IBA1+ microglia in the ipsilateral SN. **(A)** Representative effector state morphologies from segmented IBA1+ microglia. Scale bar is 10 μm. **(B)** Quantification of the average frequency of IBA1+ microglia demonstrated a microgliosis phenotype. URMC-099 reduces the frequency of microgliosis in αSyn+ cells compared to vehicle-treated rats (p = 0.0399). **(C)** Quantification of the average frequency of IBA1+ microglia demonstrating a “hyper-ramified” morphology. URMC-099 treatment reduces the frequency of hyper-ramified IBA1+ αSyn+ microglia compared to IBA1+ αSyn-microglia (p = 0.032) as well as compared to IBA1+ αSyn+ microglia in vehicle-treated rats (p = 0.0391). **(D)** Quantification of the average frequency of microglia demonstrating an “amoeboid” morphology. No differences were measured based on therapeutic regimen when comparing within cells that were either αSyn- or αSyn+, but a significant reduction in the frequency of “amoeboid” IBA1+ microglia was observed for both treatment groups when comparing αSyn+ to αSyn-(p < 0.0001). All statistical tests were performed as two-way ANOVAs with Fisher’s LSD post-hoc corrections (n = 8 per treatment) and plotted as box plots for mean and interquartile range, with whiskers spanning to min and max values.

IBA1+ cells were deemed as demonstrating microgliosis (Figure 6B) if they demonstrated expression levels (defined as IBA1 intensity normalized by the cellular volume) that were at least 2 standard deviations above the average IBA1 expression levels measured in the contralateral SN. Microglia were classified by the degree of ramification in microglia (Figure 6C) using two separate parameters – 3D ramification index (70) and fill factor (71) – that measure how a cell occupies the polyhedral space mapped by its physical extents. Specifically, these are defined by the ratio of surface areas (ramification index) and volumes (fill factor) of the cell to the encapsulating polyhedron. Microglia with higher ramification indices reflect more complex and augmented branching of their processes (70), indicative of a “primed” or early activation of a stress response (72). We used the mean values of these parameters from IBA1+ cells in the contralateral SN as an index for homeostatic ramification and defined a threshold of at least 1.5 standard deviations greater than these means to be “hyper-ramified”. Separately, microglia were also classified by their sphericity (Figure 6D), which is defined by the ratio of its surface area to volume. Values approaching the maximal limit of 1 being more spherical and amoeboid (70,71,73,74), and represent more an “amoeboid” effector state often characterized with increased motility and phagocytic activity (75).

No differences between URMC-099 and vehicle therapeutic arms were measured in the median abundance of cells demonstrating microgliosis (difference = 12.5%, p = 0.25), “hyper-ramification” (difference = 2%, p = 0.69), or “amoeboid” (difference = 2.5%, p = 0.29) effector states in IBA1+ Syn211-microglia (Figures 6B-D). Interestingly a higher fraction of Syn211-microglia are classified as “amoeboid” compared to their Syn211+ counterparts in the ipsilateral SN of both treatment cohorts (mean difference = 14%, p < 0.0001). This may reflect a population of motile activated microglia seeking debris-like αSyn for uptake and degradation.

When focusing on haSyn-burdened microglia (i.e., Syn211+ IBA1+) of the ipsilateral SN, differences between the vehicle or therapeutic cohorts were quantified. URMC-099 treatment reduced the median percentage of cells positive for microgliosis (Figure 6B) by ∼20% (p = 0.0399) as well as those classified as “hyper-ramified” (Figure 6C) by ∼50% (p = 0.0391). The aggregate populations of IBA1+ Syn211+ microglia demonstrating an “activated” morphology is reduced (difference = 9%, p 0.03) with URMC-099 treatment from 22% to 13% (Supplemental Figure S9).

### MLK3/LRRK2 co-inhibition by URMC-099 demonstrates unique transcriptomic landscape

The preservation of more homeostatic-like microglia by URMC-099 in IBA1+ Syn211+ microglia was also mirrored in the Tmem119+ microglia screened for LRRK2 activity (Supplemental Figure S10). Interestingly, URMC-099 treatment was not isolated to Tmem119+ cells with elevated LRRK2 levels (labeled LRRK2+). Rather, URMC-099 treatment had a broad influence over effector states of Tmem119+ cells in the ipsilateral SN, regardless of LRRK2 levels. This suggested that the retained ability to dynamically adopt effector states to sustain autophagic flux is likely dependent on the regulated activity of both target substrates for URMC-099, MLK3 and LRRK2. To isolate the mechanistic underpinnings of dual inhibition of MLK3/LRRK2 in microglia, we used in an *in vitro* culture model as a simplified approximation aimed at recapitulating the pre-clinical rodent model of haSyn-mediated PD. BV-2 microglia-like cells were treated with recombinant human αSyn pre-formed fibrils (PFFs, 2 μg/mL) and cotemporaneous therapeutic dosing with either vehicle or 100 nM URMC-099 (Figure 7A, Supplemental Figure S11).

**Figure 7.**
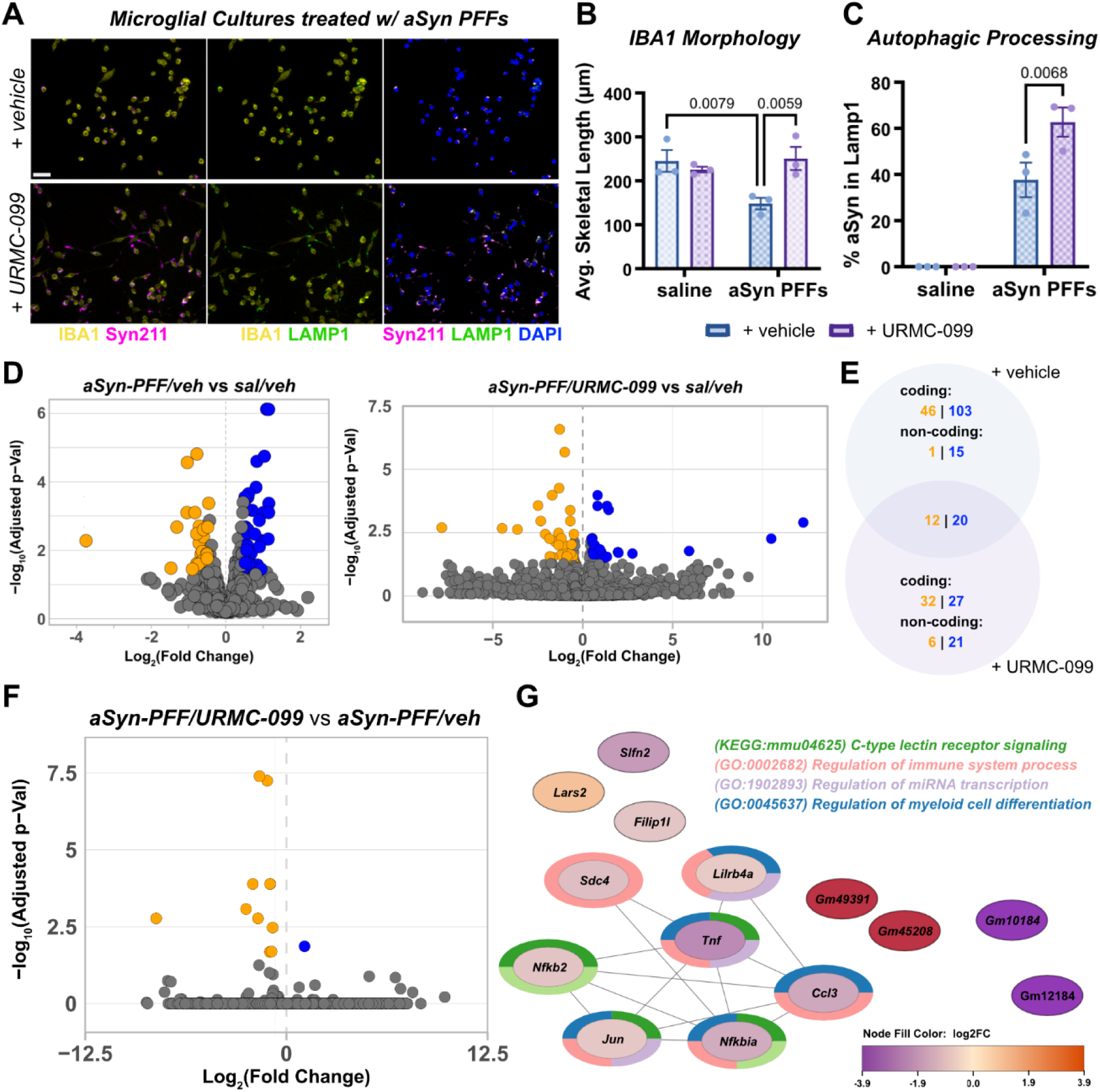
Identifying signaling modules from dual kinase inhibition using *in vitro* model for synuclein stimulated microglia. **(A)** Representative fluorescent micrographs from PFF-treated BV-2 microglia given either vehicle or URMC-099 therapy. Cells were stained for IBA1, Syn211, LAMP1, and DAPI. Scale bar is 20 μm. **(B)** Skeletonization for longest end-to-end trajectory was used to determine changes in skeletal length as a proxy for BV-2 cell morphology. PFF treatment reduced the average skeletal length ∼45% compared to negative controls (p = 0.0079) and restored by URMC-099 treatment (p = 0.0059). **(C)** Autophagic flux was evaluated via the fraction of αSyn in IBA1+ cells that were also engulfed within CD68+ structures. URMC-099 enhanced this colocalization two-fold (p = 0.0068). **(D)** Volcano plots for DEGs compared to saline negative controls in PFF-treated BV-2 cells given either vehicle or URMC-099 therapy. Orange markers reflect significantly downregulated genes. Blue markers reflect significantly upregulated genes. **(E)** Venn diagram of the distinct set of DEGs between the two therapeutic groups in PFF-treated cells when compared to the negative control. DEGs were categorized as either coding or non-coding. Orange reflects significantly downregulated genes. Blue reflects significantly upregulated genes. **(F)** Volcano plot of the DEGs directly comparing PFF-treated cells given URMC-099 to PFF-treated cells given vehicle. **(G)** Network analysis of DEGs from F with most significant signaling pathways associated with the network of interconnected genes.

After 18 hours, PFF-treated BV-2 lacking URMC-099 (PFF+/URMC-099-) demonstrated a ∼45% reduction (mean difference = 100 μm, p = 0.0079) in the average skeletal length of the IBA1+ cells from 250 μm in saline-treated negative controls to 150 μm (Figure 7B). This was reversed by URMC-099 treatment (mean difference = 100 μm, p = 0.0059) back to similar average lengths as saline-treated negative controls (mean difference = 25 μm, p = 0.39) and is reflected by the thin, extended processes the cells demonstrate (Figure 7A). These extended processes seem to correlate with the cells ability to further sequester more Syn211+ PFFs (mean difference = 600 μm^3^, p = 0.004) compared to the more amoeboid cells given vehicle therapeutic (Supplemental Figure S11). The increased sequestration of PFFs in URMC-099-treated BV-2 was complemented by a 2x increase lysosomal activity, measured as average LAMP1+ volume per cell (Supplemental Figure S11) and a corresponding 2x enhancement (mean difference = 25%, p = 0.007) in the engulfment of Syn211+ features within LAMP1+ structures, indicative of enhanced autophagy (Figure 7C). The retention of a branched morphology and enhanced autophagy in the BV-2 cells with URMC-099 was complemented by a ∼30% reduction (mean difference = 1300 a.u., p = 0.0003) of microgliosis, measured as the average expression level of IBA1 (intensity normalized by volume) in cells stimulated with PFFs compared to the IBA1 expression levels in BV-2 given vehicle therapeutic (mean expression level = 3700 a.u., Supplemental Figure S11).

We then isolated RNA for transcriptomic profiling via bulk RNA-seq to profile the unique mechanistic signature that arises from the dual inhibition of MLK3 and LRRK2 by URMC-099. As expected, stimulation with αSyn PFFs resulted in widespread expression changes in gene expression compared to the saline-treated negative controls (Figure 7D). Analysis of these differentially expressed genes (DEGs) demonstrated a highly interconnected network that resulted in the dysregulation of several pathways relevant to homeostatic cellular function and linked to PD (Supplemental Figure S12). Interestingly, URMC-099 treatment did not result in result in a DEG profile that was completely restored to that of the negative control (Figure 7D). Network analysis of these DEGs resulted in segregation into isolated, sparsely connected subnetworks that primarily downregulated transcription factor modules (Supplemental Figure S13). Approximately half of the DEGs were unable to be linked to any network or cluster. These differences in the effector signaling modules are reflected by the fact that the two treatment groups resulted in distinct DEGs when compared to the saline negative controls (Figure 7E). While direct comparison of the two therapeutic arms in only PFF-treated cells resulted in a limited number of DEGs, half of the DEGs interconnected into a network that was linked to the regulation of inflammatory and cell fate effector states (Figures 7F and 7G). Moreover, URMC-099 treatment of PFF-treated BV-2 also resulted in a distinct DEG landscape from PFF-treated BV-2 given therapeutic doses of either a LRRK2 inhibitor (Supplemental Figure S14) or MLK3 inhibitor (Supplemental Figure S15).

## Discussion

A key success metric for disease modification trials of PD is to halt the loss of neurons, both within motor circuits and outside of the basal ganglia (11,76). While advancements in clinical diagnostic methods have improved the identification of individuals potentially at risk for early PD, the disease is highly heterogeneous; thus, current therapeutic measures have not demonstrated similar widespread utility for the diverse etiologies of PD (12,20,22,23). This shortcoming may be explained by the design of many therapeutic approaches around high-risk mutations implicated in severe disease in a subset of PD patients, for example, LRRK2. Herein, we evaluate the therapeutic implications for dual inhibition of MLK3 and LRRK2 with URMC-099 and show that resilience against αSyn neuropathology involves regulating the activity profile of both kinases. Targeting of both MLK3 and LRRK2 pathways ameliorated multiple pathological features of PD. Accordingly, URMC-099 may reflect an approach to PD treatment that reflects the heterogeneous nature of the disease potential benefit for the broad spectrum of PD-associated etiologies.

To demonstrate the role of this multi-kinase pathway and investigate the therapeutic potential of URMC-099 for disease modification, we implemented a low-dose regimen of our unilateral AAV-haSyn overexpression system to specifically model progressive early PD (35). Unilateral, intranigral injection of AAV-haSyn in rats of the vehicle therapy cohort resulted in clear neurodegenerative pathology in the ipsilateral SN with accompanying lateralized motor deficits in forepaw use (Figure 1). By evaluating subregion specific immunoreactivity of TH and HuC/D in the SNpc and SNpr, we confirmed a distinct susceptibility to haSyn burden in dopaminergic neurons. In this early model, dopaminergic degeneration was primarily found to correspond to the weakening and loss of neuronal processes, but not total counts in TH+ dopaminergic neurons (Supplemental Figure S1). The dopaminergic neurodegenerative phenotype extended along the nigral-striatal axis (Supplemental Figure S6) leading to significant loss of glutamatergic synaptic densities in the striatum. This corroborates the manifestation of motor deficits that arise from dysregulation in dopamine-glutamate feedback between the nigrostriatal and corticostriatal pathways (58,59,77,78).

Rats in the therapeutic cohort were given once-daily i.p. 10 mg/kg URMC-099 for four weeks and demonstrated retention of bilateral forepaw usage (Figure 1B). Correspondingly, we measured significant attenuation of dopaminergic degeneration along the nigrostriatal axis, which we ascribed to the concomitant reduction in the levels of AAV-induced haSyn (i.e., Syn211+ features) in the SN (Figure 1C). Not only were the overall levels of haSyn lower, but the presentation of pathologic aggregates of pSer129-αSyn was reduced with URMC-099 treatment (Figures 3A-C). Intriguingly, no differences were observed in the formation of proteinase K resistant synuclein deposits between the vehicle and URMC-099 cohorts (Supplemental Figure S16). We hypothesize that this may reflect a stable, “benign” species of αSyn aggregates that form as a method of protecting against oxidative stress, like that observed for amyloid (79). Thus, regulation of MLK3 and LRRK2 activity is not expected to increase the breakdown of these insoluble, proteinase K resistant aggregates. Based on these results, we concluded that once-daily i.p. of 10 mg/kg URMC-099 effectively limited the progression of PD-associated pathology and prevented the progression of motor deficits in our pre-clinical model of progressive PD.

We then confirmed that the therapeutic activity of URMC-099 was indeed through engagement of its target substrates – MLK3 and LRRK2 (Figure 2). Inflammatory stress in PD clinical phenotypes involves multiple kinase signaling pathways relevant to neuronal and glial function and fate (24,80,81), with dysfunctional MLK3 and LRRK2 overactivity extensively implicated across a broad spectrum of disease etiologies. A key indicator of elevated MLK3 and LRRK2 activity is an extensive phosphorylation burden of the two kinases (i.e., pMLK3 and pLRRK2 expression levels). We find that in the early progressive model, pMLK3 and pLRRK2 burden is localized to the lesion-bearing, ipsilateral SN at 6 weeks post-induction. The phosphorylation burden of MLK3 and LRRK2 is greatly alleviated with once-daily URMC-099, though there was a marked difference in the magnitude of reduction pMLK3 compared to pLRRK2. We attribute this to the fact that MLK3 regulation of inflammatory stress is more ubiquitous across cell types compared to LRRK2 (33,82). This rationalization is corroborated by not only improved cellular integrity of nigral neurons (Figures 1D and 1E and Figure 4) but also reduced inflammatory activation in nigral astrocytes and microglia (Figures 3D and 3E).

Separately, dysregulated LRRK2 activity manifests distinct pathologies between neurons and microglia. In neurons, we posit that elevated LRRK2 activity most negatively impacts neuronal health at synapses by dysregulating vesicular trafficking activity (83,84). Additionally, while aberrant LRRK2 signaling is broadly linked to dysregulated autophagic flux, we posit that this pathologic function has a greater impact on microglia and their ability effectively ameliorate the neurodegenerative stress (45,81,85).

In nigral neurons, we found that improved neuronal health with URMC-099 treatment directly correlated with restored levels of VAMP expression and a reduction in the formation of larger aggregates of pathologic αSyn along neuronal processes (Figure 4A-C). LRRK2 dysfunction has demonstrated regulatory function over VAMP4 and VAMP7 in the context of αSyn-mediated stress and trafficking, with downstream consequences in the trafficking of VAMP2 to pre-synaptic terminals (86,87). Consequentially, VAMP2-related chaperone function has been implicated in the regulation of αSyn association into biomolecular condensates and is positively correlated with reduced aggregation potential of αSyn. Thus, the reduced levels of pre-synaptic VAMP2 from the lack of LRRK2 regulation from URMC-099 therapy is likely linked to the increase in pathologic αSyn accumulation and resultant loss of neuronal integrity. This is demonstrated by the rescue of MAP2 integrity along neuronal processes (Figure 4A). We also observed bright, dense MAP2 expression patterns that are colocalized with hαSyn inclusions in the soma of neurons in the SN from the vehicle cohort (Supplemental Figure 5); these are reduced in frequency in the therapeutic cohort. We attribute this observation to abnormal patterns in MAP2 localization, such as the formation of fibrous structures, with nuclear/somatic αSyn inclusions that have been implicated as part of cellular Lewy body pathology (88).

From single-cell segmentation analysis of IBA1+ and Tmem119+ microglia, we measured a clear reduction in the phosphorylation burden on LRRK2 in microglia, but this did not correlate with an increased percentage in the number of microglia with Syn211+ haSyn engulfed with CD68-labeled lysosomal structures (Figure 5). This discrepancy with the clear reductions in total and pathologic αSyn burden (Figure 1C and Figures 3A-C) was linked to a need for microglia to efficiently perform autophagy but also stay resilient to chronic proteostatic stress. Specifically, we propose that URMC-099-mediated enhancement of autophagic flux as a product of both regulation of LRRK2-related autophagic activity and MLK3-related inflammatory stress.

By performing morphological analysis of microglia effector states (Figure 6 and Supplemental Figures S9 and S10), we identified a significant retention of homeostatic-like αSyn-burdened microglia with URMC-099 treatment that was not strongly correlated to the degree of LRRK2 activity in a cell. Specifically, we classified microglia into three established morphologic effector states: ramified (homeostatic, surveilling and phagocytic), hyper-ramified (stressed), and amoeboid (migratory and phagocytic). Interestingly, in both therapeutic arms we found a lower fraction of IBA1+ cells displaying an “activated” morphology if they were Syn211+ (Supplemental Figure S9); this was further reduced for Syn211+ microglia in the URMC-099 therapy arm. Disaggregation of the activated morphologies into their specific classifications identified a significant transition away from amoeboid morphologies in Syn211+ cells, possibly reflecting the transition into a less phagocytic effector state upon engulfment of αSyn to utilize intracellular machinery for processing and degradation. Microglia from rats given the vehicle therapeutic demonstrated a greater fraction of hyper-ramified cells and cells demonstrating gliosis, both of which URMC-099 significantly reduced and suggests a mechanistic push towards a more homeostatic or surveilling phenotypes in the microglia.

A sustained population of activated microglia may lead to aberrant phagocytic activity that can lead to unwanted pruning of synaptic structures or conversion of stable, “benign” insoluble aggregates into neurotoxic forms (89,90). Conversely, a greater population of homeostatic-like microglia likely implicates preserved dynamicity of microglia to transition between effector states to more effectively maintain a neuroprotective environment. While we did not find a reduction of proteinase K resistant Syn211+ aggregates in the SN between therapeutic groups (Supplemental Figure 16), we did observe a reduction in the frequency of Homer1+ glutamatergic synapses surveilled by microglia along the nigral-striatal pathway (Supplemental Figure S17). Combined with the overall reduction in pathologic αSyn levels (Figure 3) in the ipsilateral SN and retention of Homer1+ synapses in the striatum (Supplemental Figure 6), our data suggests a neuroprotective effector state in microglia that is promoted by dual inhibition of MLK3 and LRRK2 by URMC-099. This we attribute back to the preservation of dynamicity in the microglia to both transition into and out of different effector states, mediated by regulation of autophagic flux by LRRK2 and inflammatory stress by MLK3.

Lastly, we tested our proposed dual inhibition of MLK3/LRRK2 regulatory module as a resilience mechanism in microglia to retain their ability to dynamically transition between effector states to enhance overall autophagic flux. We evaluated differences in the transcriptomic signature of microglia-like BV-2 cells in a model for autophagic activity following treatment with αSyn PFFs and co-treatment with either a MLK3 inhibitor, LRRK2 inhibitor, or URMC-099 (Figure 7 and Supplemental Figures S11-15). URMC-099 treatment only produced a subset of DEGs that overlapped with the DEGs from sole inhibition of either MLK3 or LRRK2. Downstream functional network analysis and ontology mapping further demonstrated distinct effector signaling modules from dual inhibition by URMC-099 compared to single target inhibition of either MLK3 or LRRK2.

### Conclusions

In aggregate, our study provides a convergence of evidence to suggest a unique therapeutic mechanism from dual kinase inhibition of MLK3 and LRRK2 by URMC-099 relevant to PD.

## Materials and Methods

A full list of materials and extended methods are included in the supplementary document.

### AAV Production

The self-complementary AAV genome encoding human α-syn (pscAAV-CBh-aSyn-WPRE3-enSV40pA) was a gift from Michael J Fox Foundation (Addgene plasmid: 194244; http://n2t.net/addgene:194244; RRID: Addgene_194244). The genome was packaged into AAV2/5 as described previously using a double-transfection protocol of 293T cells using the genome plasmid and pXYZ5 containing AAV5 cap and helper functions. Vector was purified using an iodixanol gradient, dialyzed using modified PBS, and titered using ddPCR (91,92). The vector was diluted to the working titer (1.7 x 10^12^ vector genomes (vg)/ml) using balanced salt solution (Alcon). This titer was intended to yield mild to moderate pathology with minimal neurodegeneration within the experimental time window as described (36).

### Animals

Young adult male and female Sprague-Dawley rats (3-4 months old) were obtained from Charles River Laboratories, and all procedures were performed in accordance with the Barrow Neurological Institute Institutional Animal Care and Use Committee (IACUC). Animals were group housed with 2-4 animals per cage, maintained on a controlled light cycle (12-hour light/dark) and temperature environment (22 °C) with food and water available *ad libitum*.

### Stereotaxic Delivery of AAV

Surgery was performed under isoflurane anesthesia (1.7% - 2.0%). Animals were fitted into a stereotaxic frame and all animals received a 3 µl unilateral injection of AAV 2/5 pscAAV-CBh-aSyn-WPRE3-enSV40pA into the SN using standard methodology described elsewhere (93). Briefly, a glass capillary coated with SigmaCote was fitted to a 10 µl Hamilton Gas Tight syringe attached to a motorized injector (Quintessential Stereotaxic Injector, Stoelting). Coordinates relative to Bregma were AP - 5.3 mm, ML +1.8 mm, DV - 7.2 mm (from dura). Virus was injected at a rate of 0.4 ul/min and needle was held in place for an additional 5 minutes before retracting.

### URMC-099 Administration

Once-daily administration of URMC-099 began 14 days after stereotaxic vector delivery. URMC-099 was dissolved as a 10 mg/mL working solution with 5% v/v DMSO (Sigma), 40% v/v PEG-400 (Millipore) in saline (0.9% NaCl). Animals with administered (i.p.) either 10 mg/kg of URMC-099 or an equivalent volume of vehicle control.

### Cylinder testing

The day before sacrifice animals underwent cylinder testing for asymmetric forepaw use (94). Briefly, animals were placed in a clear plexiglass cylinder and video recorded until the animal performed 20 weight-bearing paw placement on the cylinder, or until 5 minutes had elapsed. A blinded observer quantified left, right, and simultaneous paw placement from recorded videos. The left and right paws were then classified as contralateral or ipsilateral with respect to the location of the unilateral brain lesion associated with AAV-haSyn injection. Data is presented as the % of total paw usage that involves the contralateral forepaw, with symmetric forepaw usage defined as 50% use of the contralateral forepaw. This is quantified by the following equations:

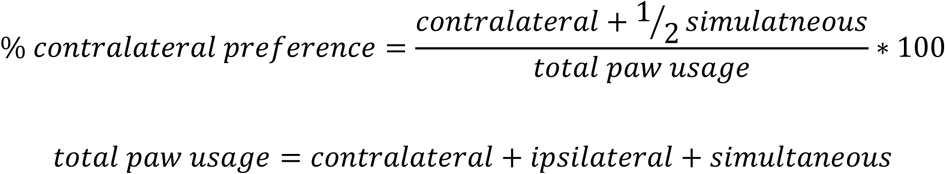

### Euthanasia

Six weeks following vector delivery animals were sacrificed using methodology in accordance with the recommendations of the American Veterinary Medical Association and as approved by the Barrow Neurological Institute IACUC. Animals were deeply anesthetized with 60 mg/kg pentobarbital (i.p.) and thereafter transcardially perfused with Tyrode’s solution followed by 4% ice-cold paraformaldehyde, postfixed in 4% paraformaldehyde, and thereafter placed in 30% v/v sucrose in 0.1 M phosphate buffer for cryoprotection. Brains were thereafter cut coronally into 40 µm thick sections using a sliding stage microtome. Sections for each brain were serially distributed into 6 groups and stored in cryoprotectant at −20 °C (30% v/v glycerol, 30% PEG300, 20% 0.1 M phosphate buffer, and 20% dH2O) at −20 °C) until further use. Moreover, a subset of tissue was shipped to University of Rochester Medical Center (URMC) for processing.

### Brain Histology

#### Immunohistochemistry

All immunohistochemistry was performed on free-floating sections as previously described (95). Brightfield sections were washed thrice in 1x PBS (30 minutes) quenched with 0.3% hydrogen peroxide, followed by blocking in the appropriate serum, 2 hrs at room temperature (RT). Following the PBS wash, tissue for immunofluorescence and near-infrared imaging was washed in 100mM of glycine in 1x PBS (30 minutes) to remove tissue and fixation autofluorescence. Sections were then washed (3 times; 10 minutes) in 1x PBS to remove excess glycine, before incubation in antigen unmasking solution (Vector Laboratories; H3300) at −20 °C to improve antigen-antibody binding and washed (3 times; 10 minutes) in 1x PBS.

Brightfield sections were thereafter incubated with the primary antibody (TH, HuC/D) at RT overnight, followed by the secondary antibody for 2 hr at RT. Sections were incubated using the Vectastain ABC kit (Vector Laboratories) per developer instructions and developed using 3,3′ diaminobenzidine and 0.03% hydrogen peroxide in tris buffer. Sections were mounted on subbed slides, dehydrated using increasing concentrations of ethanol followed by xylene and coverslipped with Cytoseal™ (Fisher Scientific).

Immunofluorescent (IF) sections were incubated in their respective primary antibodies (LRRK2, pLRRK2, pMLK3, IBA1, CD68, αSyn, GFAP) in staining buffer (3% normal goat serum + 1.5% bovine serum albumin in 1x PBS with 1.8% NaCl and 0.5% Triton X-100) for 3-days at RT. Following incubation in primary antibodies, sections were washed thrice in 1x PBS + 1.8% NaCl (30 min). Sections were incubated in their respective Alexa-fluor conjugated goat secondary antibodies (ThermoFisher; 1:500) in staining buffer overnight. Following secondary antibody staining, sections were washed thrice in 1x PBS + 1.8% NaCl (30 minutes each) and mounted on microscope slides (VWR) and coverslipped with ProLong Diamond Antifade Mounting Media with or without DAPI (ThermoFisher) and #1.5 coverslips (VWR). Slides were cured overnight prior to imaging.

#### Proteinase K digestion

Proteinase K digestion was performed as described previously (96). Briefly, tissue was incubated with Proteinase K (ThermoFisher) at 50 μg/mL in TBS-T (10 mM Tris-HCl, pH 7.6; 100 mM NaCl; 0.1% Tween-20) for 40 min at 50 °C. Sections were thereafter immunolabeled using the anti-α-synuclein antibody Syn211, followed by a biotin-SP-conjugated goat anti-mouse IgG secondary antibody. High-resolution brightfield photomicrographs were subsequently acquired using a ZEISS Axioscan.Z1 system.

### Imaging

#### Brightfield images

DAB-stained images for brightfield imaging were captured on a Nikon Eclipse Ni microscope and processed using IS-Elements software (Nikon Instruments Inc.)

#### AI-based enumeration of TH+ cell bodies

Slides for AI-based enumeration of TH+ cells were captured using a ZEISS Axioscan (ZEISS Group; Oberkochen, Germany). Serial images consisted of a series of image tiles acquired across the X-Y plane of the SN. In addition, each tile consisted of a stack of images acquired with a 20X objective across 13 µm on the Z-axis with a 0.8 µM step size. Acquired photomicrographs were processed and stitched using the ZEN® advanced software, producing a single high-resolution digital image per each scan which were uploaded to AIforia (AIforia technologies, Finland) for subsequent enumeration of TH + cells (97).

#### Quantitative near-infrared imaging

Whole brain mounted slices for immunofluorescent imaging of TH in the striatum was acquired using an Odyssey CLx near-infrared imaging system (LI-COR Biosciences) on 40-µm brain sections in the 700-nm channel using an intensity setting of 2.5, 169-µm resolution, high-quality acquisition, and a 1.0-mm focus offset.

#### Volumetric z-stack IF Microscopy

Volumetric z-stack images for IF microscopy data was acquired using an Olympus BX-51 microscope in “grid-confocal” configuration using an OptiGrid structured illumination optical element (Qioptiq) in the excitation path as previously reported (98). A Prior Lumen200 illumination source equipped with a Hg lamp (Prior; cat: LM200B1-A) is coupled into the microscope via a liquid light coupler (Prior; cat: LM587). Fluorescent emission from the immunostained samples was detected using a Hamamatsu ORCA-ER scientific camera. The following pairs of excitation and emission filters were used to selectively illuminate and detect biomolecular targets immunolabeled with DAPI, AlexaFluor488, AlexaFluor568, and AlexaFluor647, respectively: 350 nm / DAPI (Semrock; cat: FF02-447/60-25); 405 nm / FITC (Semrock; cat: FF01-524/24-25); 488 nm / TRITC (Semrock; cat: FF01-593/40-25); 568 nm / Cy5 (Semrock; cat: FF01-692/40-25). Volocity 3DM software (Quorum Technologies) was used for hardware integration and control to acquire all images. Synaptic punctae in the striatum were imaged at 60x magnification (0.5 μm z-step sizes), while structural detail of neurons in the substantia nigra were acquired at 100x magnification (0.3 μm z-step sizes). Other markers were imaged at 20x magnification at (1 μm z-step sizes). The following infinity-corrected Olympus objectives were used: 20x (UPlanApo 0.70 NA), 60x (UPlanSApo 1.35 NA oil); 100x (UPlanSApo 1.40 NA oil).

### Image Analyses

#### AI-based enumeration of nigral TH+ cells

AI-based enumeration was performed using the AIforia platform (AIforia Technologies; Helsinki, Finland) as previously described (97). Briefly, this platform uses an AI deep convolutional neural network (CNN) learning to facilitate quantitative histological image analysis. The region of interest (SNpc) was outlined and the CNN was used to enumerate the total number of TH+ cells. We previously validated this approach by performing standard TH stereology on the same sections and fit the results with a linear regression analysis that showed high concordance between the methodologies (95). In this study, we used a smaller sample set (3 animals from each cohort) to confirm alignment with our previous studies (R^2^ = 0.91, p = 0.003).

#### Li-Cor based densitometry

To quantify striatal TH immunoreactivity following URMC-099 treatment, 40-µm brain sections were immunolabeled with an anti-TH primary antibody. A minimum of 10 sections per animal, corresponding to every sixth section distributed throughout the rostrocaudal extent of the striatum, were included in the analysis. The entire anatomical boundaries of the striatum were outlined based on Paxinos and Watson rat brain atlas. A cortical region of interest on the naïve hemisphere was used to define background signal. For each hemisphere, the total TH signal from all analyzed sections was summed and divided by the summed striatal area to obtain TH signal per unit area. TH immunoreactivity in the lesioned hemisphere is expressed as a percentage of the corresponding naïve hemisphere.

#### Proteinase K densitometry

Quantification of staining intensity was performed using FIJI/ImageJ software (National Institutes of Health). Prior to analysis, all images were converted to 8-bit grayscale and inverted to optimize pixel intensity mapping, followed by background subtraction using a rolling ball radius of 50 pixels to minimize non-specific signal. Quantitative measurement parameters, including area, mean gray value, minimum and maximum gray values, and threshold limits, were configured and uniformly applied across all images to ensure consistent analysis. The SNpc was manually delineated with reference to the Paxinos and Watson rat brain atlas. A minimum of two sections per rat were included in the analysis. Once regions of interest were defined, densitometric analysis was performed using the Measure function in FIJI/ImageJ. Mean gray value was used as the primary metric for quantifying staining intensity, and the resulting values were exported for statistical analysis. Data are reported in arbitrary units as the ratio of staining intensity between hemispheres.

#### DAB-stained brightfield images

Brightfield images of TH and HuC/D staining were analyzed using ImageJ (NIH). The general workflow is as follows. Cropped brightfield images of the midbrain from each hemisphere were loaded into ImageJ and aligned to cropped reference images from the Waxholm brain atlas of the same coronal depth. The atlas reference images were labeled for the SNpc and SNpr in different colors (default from the web tool). Alignment was performed using the BigWarp plugin (v 9.3.0) with the brain atlas as the moving image and the DAB-stained brightfield image as the target image (37). A binary mask images were generated for the brightfield image, as well as individual area maps for the SNpc and SNpr that were extracted from the color-coded atlas using the “Color Threshold” tool. The binary “AND” operation was used to generate masks of immunoreactive signal specific to the SNpc and SNpr. These sub-region specific masks were then normalized to a max signal intensity of 1, and multiplied against the original brightfield image to extract measurements on “area” “integrated density” and “area fraction” using the “Measure” tool on ImageJ. Extracted values were exported as a .csv for data cleaning, followed by imported into GraphPad Prism for plotting and statistical analysis.

#### Immunofluorescence

Semi-quantitative data was extracted from volumetric IF z-stack images using Volocity 3DM Quantitation Software (Quorum Technologies). The general scheme was as follows. A fine noise filter was used on all images before applying a set of measurement protocols. The “Find Objects” function was used to identify fluorescent objects above a certain intensity threshold defined by n standard deviations above the average voxel intensity of a given channel, where n ranged from 1.5 to 3 standard deviations depending on the skew of the intensity distributions in each channel. Colocalization of immunoreactive signals regardless of cellular boundaries was determined using the “Intersect” function. All processed objects for individual channels were exported to a .csv that was imported into MATLAB for data cleaning, filtering, and calculation of additional semi-quantitative parameters (i.e., intensity / volume, relative %, morphological parameters, and population-based statistics). Summarized data values were imported into GraphPad Prism for plotting and statistical analysis. Data was plotted either as the mean with 95% confidence intervals or as box plots (mean + interquartile ranges) with whiskers that spanned the min and max range. Full details are provided in the supplemental text.

### Cell culture model to recapitulate haSyn-mediated PD mechanisms in microglia Microglia-like BV-2 Cell Culture

Microglia cultures were comprised of an immortalized murine microglia-like cell line (BV-2; RRID: CVCL_0182) grown on tissue-culture substrates coated with 100 μg/mL of poly-D-Lysine (PDL) in ddH_2_O and maintained as previously reported (99,100). Cells were maintained in culture medium comprised of Dulbecco’s Modified Eagle Medium (DMEM, ThermoFisher) supplemented with 10% v/v fetal bovine serum (FBS, Atlas Biologicals), 1% v/v GlutaMax (ThermoFisher), and 1% v/v penicillin-streptomycin (ThermoFisher) at 37 °C and 5 % CO_2_. Cells were passaged when confluency was between 75% and 85% confluency using 0.25% trypsin to dissociate them for the surface of the culture flask. Cells used in experiments were from passage numbers p12 to p16. Cells used for transcriptomics were cultured on PDL-coated 12-well plates (Corning) at ∼85k cells/well, while cells used for ICC were cultured on plasma-cleaned (20 minute) and PDL-coated coverglass (12mm, Neuvitro) in 24-well plates (Corning) at ∼40k cells/well. Before treatment, the cultures were incubated for 1 hour in a reduced serum condition (FBS reduced to 1% v/v). This was followed by overnight (16 - 20 hours) incubation at 37 °C and 5 % CO_2_ with vehicle (0.1% v/v saline + 0.1% v/v DMSO), aSyn PFFs (2 ug/mL + 0.1% v/v DMSO), URMC-099 (100 nM + 0.1% v/v saline), or both aSyn PFFs (2 ug/mL) and URMC-099 (100 nM) in reduced serum medium. The cells were then either fixed for ICC or collected for RNA extraction. aSyn PFFs (StressMarq) were sonicated at power level 7 (VWR Model 50D) for 5 minutes at 20 °C prior to use.

### Immunocytochemistry

BV-2 microglia cultures on coverslips were stained using an indirect immunofluorescent labeling protocol as previously reported (99,100). Cultures were briefly rinsed with ice-cold DPBS, followed by fixation in 4 % paraformaldehyde (PFA) diluted in a 1x PBS solution for 12 minutes at room temperature and gentle rocking. The fixative is then removed and quenched with 100 mM glycine (BioRad) in 1x PBS solution for 5 minutes at RT, followed by two washes with 1x PBS at the same conditions. The cells were then permeabilized using 0.25 % Triton X-100 (Millipore Sigma) in ddH2O 1x PBS for 15 minutes at room temperature and gentle rocking, followed by one wash with 1x PBS in the same conditions. The cells were then blocked using 10 % BSA (Millipore Sigma) for 1 hr at RT and gentle rocking. Primary antibodies (IBA1, Syn211, LAMP1) diluted in staining buffer (3 % BSA + 0.05% Triton X-100 in 1x PBS) were incubated at 4 °C overnight with gentle rocking, followed by three washes with 1x PBS (5 min). AlexaFluor-conjugated secondary antibodies (ThermoFisher; 1:750) were diluted in the same staining buffer and allowed to incubate for 1 hour at RT with gentle rocking. The cells were then washed three times with PBS-T (1x PBS + 0.1% Tween-20) for 5 minutes each. After staining, the coverslips were mounted onto cover glass using ProLong^TM^ Diamond Antifade Mountant with DAPI.

### Imaging and Analysis

Volumetric z-stack images were acquired and analyzed using Volocity 3DM Acquisition and Quantitation software, respectively. The same “grid-confocal” microscope system and analysis workflow as the immunohistochemical IF images was used here.

### Transcriptomics

#### RNA Extraction

RNA was isolated from BV-2 microglia using a *Quick*-RNA Miniprep Kit (Zymogen), following the manufacturer recommended protocol with no modifications. After isolation, the purity (A260/A280 and A260/A230) and concentration (A260) of the extracted RNA was determined using a NanoDrop^TM^ Spectrophotometer (ThermoFisher). Only total RNA of acceptable quality (A260/A280 > 1.8, A260/A230 > 1.5) were then stored at −80 °C.

#### RNA-seq library generation and mapping

Input RNA was subjected to polyA selection using NEBNext Poly(A) mRNA Magnetic Isolation Module (NEB). 2 µg of RNA from each sample was used for kit purification and were eluted in 10 mM Tris HCl pH7.5 before proceeding to library generation. RNA-seq libraries were generated (3 independent biological replicates per condition) using the Click-seq library preparation method with a 1:35 azido-nucleotide ratio (101). Libraries were sequenced using a single-end 150 bp cycle run on an Illumina NovaSeq 6000.

Raw reads were generated from the Illumina basecalls by demultiplexing using bcl2fastq (v 2.20.0) followed by quality filtering and adapter removal using fastp (v 0.23.1). Processed reads were then mapped to the GRCm39 reference genome using STAR.

#### Analysis of RNA-seq read counts

Gene level read quantification was derived using the featureCounts package with a GTF annotation file (GRCm39) (102). Differential expression analysis was performed using DESeq2-1.34.0 with a P-value threshold of 0.05 and log2FC of 0.5. Data was analyzed using HyperGen genomics analytics software, a Hyperion application, developed by the Wilmot Cancer Institute’s Technology and Innovation Group automating open-source and customized bioinformatics tools from the University of Rochester Genomics Research Center. Gene ontology and pathway analyses were performed on Cytoscape with the ClueGO, stringApp, ReactomeFI, clusterMaker2, and EnrichmentMap plugins (103–108). Full details on read processing, mapping, and quantification are provided in the supplemental text.

### Statistics

All statistics reported were computed using GraphPad Prism 11.0.0. Analyses involving only two groups were performed as two-tailed non-parametric Mann-Whitney tests. These include experimental readouts with comparisons that only focused on lateralized differences due to AAV-haSyn injection, regardless of therapeutic cohort, and those that only focused on differences due to therapy in the lesion-bearing hemisphere. Analyses focused on the interactions of AAV-induced haSyn expression and dual inhibition of MLK3/LRRK2 by once-daily URMC-099 therapy were performed as two-way ANOVAs with Fisher’s LSD post-hoc corrections to extract direct group contrasts. For RNA-seq, statistical significance was determined using Mann-Whitney pairwise tests unless otherwise noted. Benjamini-Hochberg False-discovery rate corrections are performed by DESeq2 during DEG analysis.

Full details regarding sample sizes, significance tests, and plotted values are included with the accompanying legend for each figure.

## Declarations

### Ethics approval and consent to participate

All animals were handled in alignment with approved protocols and guidelines in accordance with Barrow Neurological Institute IACUC.

### Availability of data and materials

All data required to reproduce the figures reported in the main text and supplemental document will be provided as supplemental data files upon publication. Cleaned data and their respective analysis protocols to reproduce the quantifications and results are also available in a data repository stored on OSF.io accessible at the following link: https://osf.io/vgzj3/overview?view_only=08e10db40c124957980664995904c1bc.

RNA-seq datasets will be made available freely available on the NIH GEO website under accession number.

## List of Abbreviations

AAV: adeno-associated virus
CNN: convolutional neuronal network
DAB: diaminobenzidine
DEG: differentially expressed genes
DMEM: Dulbecco’s Modified Eagle Medium
FBS: fetal bovine serum
GFAP: glial fibrillary acidic protein
haSyn: human alpha-synuclein
HuC/D: Hu antigens C/D
i.p.: intraperitoneal
IBA1: ionized calcium-binding adaptor molecule 1
ICC: immunocytochemistry
IF: immunofluorescence
IHC: immunohistochemistry
LAMP1: lysosome associated membrane protein 1
LRRK2: leucine rich repeat kinase 2
MAP2: microtubule-associated protein 2
MLK3: mixed lineage kinase 3
PD: Parkinson’s Disease
PFA: paraformaldehyde
PFF: pre-formed fibril
pLRRK2: phosphorylated LRRK2
pMLK3: phosphorylated MLK3
pSer129: phospho-Ser129 alpha-synuclein
SD: Sprague-Dawley
SN: substantia nigra
SNpc: substantia nigra pars compacta
SNpr: substantia nigra pars reticulata
TH: tyrosine hydroxylase
VAMP: vesicle-associated membrane protein
αSyn: alpha-synuclein

