## Supplemental Document for "Dual kinase inhibition provides early clues to restoration of neuroimmune homeostasis in an AAV-haSyn model of Parkinson’s disease"

[#] *Co-first authors*

[*] *Corresponding Authors*

### 1. Extended Methods

#### Image Analyses

##### *Immunofluorescence*

After finding IBA1+ and Tmem119+ immunoreactive signal in the SN using the “Find Objects” function, individual IBA1+ or Tmem119+ microglia were segmented using the “Separate Touching Objects” with a volume cutoff that reproducibly processed larger objects into 1 or fewer microglia. Volume and shape filters were then used to remove segmented objects that were fragments of microglia. Other protein markers imaged with microglia were also segmented based on the IBA1+ or Tmem119+ cellular boundaries and assigned as belonging to the microglia they overlap with. This was done using the “Compartmentalize” function. Extracted data regarding segmented microglia included both the surface area, volume, and intensity of individual IBA1+ or Tmem119+ cells. The coordinates of the bounding polyhedron encompassing individual microglia was also extracted.

Homer1+ synaptic puncta in the striatum were identified using Volocity 3DM Quantitation via a double-positive screen for fluorescent objects that were above the “Find Objects” threshold as identified as local maxima via the “Find Spots” function. These “colocalized” positive voxel objects were then labeled as synaptic puncta for subsequent analyses. IBA1+ microglia in the striatum were segmented using the same protocol as above, and the number of Homer1+ synaptic puncta in contact with individual microglia was assigned using the same “Compartmentalize” function.

Export of data extracted immunoreactive objects identified using the “Find Objects” function contained information regarding their volume and intensity. The exported quantifications were then imported into MATLAB for data cleaning and processing. Computations included simple analyses such as intensity / volume, as well as more complex morphological analyses that calculated the shape factor and fill factor of individual microglia, followed by population estimates that quantified the median and distributions of segmented cells based on these morphological parameters. Exact scripts and exported data structures containing both cleaned and final computed values are provided in the OSF.io data repository.

#### Transcriptomics

##### *RNA-seq read processing, mapping, and analysis*

Raw reads generated from the Illumina basecalls were demultiplexed using bcl2fastq version 2.20.0. Quality filtering and adapter removal are performed using FastP version 0.23.1 with the following parameters: "--length_required 35 --cut_front_window_size 1 --cut_front_mean_quality 13 --cut_front --cut_tail_window_size 1 --cut_tail_mean_quality 13 --cut_tail -y –r" (97). Processed/cleaned reads were then mapped to the GRCm39 reference using STAR_2.7.9a with the following parameters: "—twopass Mode Basic --runMode alignReads --outSAMtype BAM Unsorted – outSAMstrandField intronMotif --outFilterIntronMotifs RemoveNoncanonical – outReadsUnmapped Fastx" (98,99). Gene level read quantification was derived using the subread-2.0.1 package (featureCounts) with a GTF annotation file (GRCm39) and the following parameters for stranded RNA libraries "-s 2 -t exon -g gene_name" (100). Differential expression analysis was performed using DESeq2-1.34.0 with a P-value threshold of 0.05. Data was analyzed using HyperGen genomics analytics software, a Hyperion application, developed by the Wilmot Cancer Institute’s Technology and Innovation Group automating open-source and customized bioinformatics tools from the University of Rochester Genomics Research Center.

#### 2. Tables of Materials and Reagents

| Table 1: Biological Specimens | |
|---|---|
| Sprague Dawley Rats | Inotiv |
| BV-2 Cells | Gift of Dr. Sanjay B. Maggirwar |

| Table 2: Consumables | | |
|---|---|---|
| **Item Name** | **Vendor** | **Catalog** |
| Cell culture plates (12-well and 24-well formats) | Corning | 3512 and 3524 |
| 12-mm coverglass | Neuvitro | GG-12-15H |

| Table 3: Reagents | | |
|---|---|---|
| **Reagent Name** | **Vendor** | **Catalog** |
| Dulbecco's Modified Eagle Medium (DMEM) | ThermoFisher | 10313039 |
| Fetal Bovine Serum (FBS) | Atlas Biologicals | F-0500-D |
| GlutaMax Supplement | ThermoFisher | 35050061 |
| Penicillin-Streptomycin | ThermoFisher | 15140122 |
| 0.25% Trypsin | ThermoFisher | 15050-065 |
| aSyn PFF Type 1 | StressMarq | SPR-322 |
| Proteinase K | ThermoFisher | 25530015 |
| ProLong™ Diamond Antifade Mountant with DAPI | ThermoFisher | P36962 |
| Prolong Diamond Antifade Reagent | ThermoFisher | P36961 |
| Glycine | BioRad | 1610718 |
| Bovine Serum Albumin | Sigma | A3294 |
| Tween 20 | Sigma | P1379 |
| *Quick*-RNA Miniprep Kit | Zymo Research | R1055 |
| NEBNext Poly(A) mRNA Magnetic Isolation Module | New England Biolab | E7490S |
| Triton X-100 | Sigma | T9284 |
| URMC-099 | Gelbard Lab | |
| DMSO | Sigma | D1435 |
| PEG-400 | Sigma | PX1286B-2 |
| Poly-D-Lysine | Sigma | P1149 |

| Table 4: Antibodies | | | |
|---|---|---|---|
| **Antibody *(host)*** | **Associated Figure(s)** | **Vendor *(Catalog)*** | **Dilution** |
| Syn211 *(mouse; IgG1)* | 1C, 3B/C, 4A/C, 5D, 7A<br>S5, S7, S8, S11, S17 | Invitrogen *(AHB0261)* | 1:500<br>or 1000 (for S17) |
| TH *(rabbit; IgG)* | 1D | Millipore *(AB152)* | 1:4000 |
| HuC/D | 1E | | |
| pMLK3 *(rabbit, IgG)* (Thr277, Ser281) | 2B | Invitrogen *(PA5-105817)* | 1:200 |
| pLRRK2 *(rabbit, IgG)* (Ser935) | 2C, 5A | Invitrogen *(701066)* | 1:200 |
| pSer129 *(rabbit, IgG)* | 3A/C, 4A/C | Abcam *(ab51253)* | 1:100 |
| GFAP *(chicken)* | 3D<br>S3 | Neuromics *(CH22102)* | 1:1000 |
| IBA1 *(rabbit, IgG)* | 3E, 5D<br>S4, S8, S16 | Wako *(PTR2404)* | 1:500 |
| IBA1 *(chicken; IgY)* | 7A<br>S11 | Neuromics *(CH22134)* | 1:4000 |
| MAP2 *(chicken; IgY)* | 4A/C<br>S5, S7 | Encor *(NC0388389)* | 1:5000 |
| VAMP1/2/3 *(guinea pig)* | 4A/C | Synaptic Systems *(104 104)* | 1:500 |
| Tmem119 *(guinea pig)* | 5A | Synaptic Systems *(400 004)* | 1:500 |
| LRRK2 *(mouse; IgG1)* | 5A | Novus Bio *(NBP2-42175)* | 1:200 |
| CD68 *(rat; IgG2a)* | 5D | BioRad *(MCA1957GA)* | 1:500 |
| LAMP1 *(rabbit, IgG)* | 7A | Cell Signaling *(9091S)* | 1:250 |
| Homer1 *(chicken)* | S6, S16 | Synaptic Systems *(160 006)* | 1:500 |
| Anti-mouse 2$^{o}$ Peroxidase conjugated *(goat)* | S17 | Sigma *(AP124P)* | 1:500 |

##### 3. Supplemental Figures

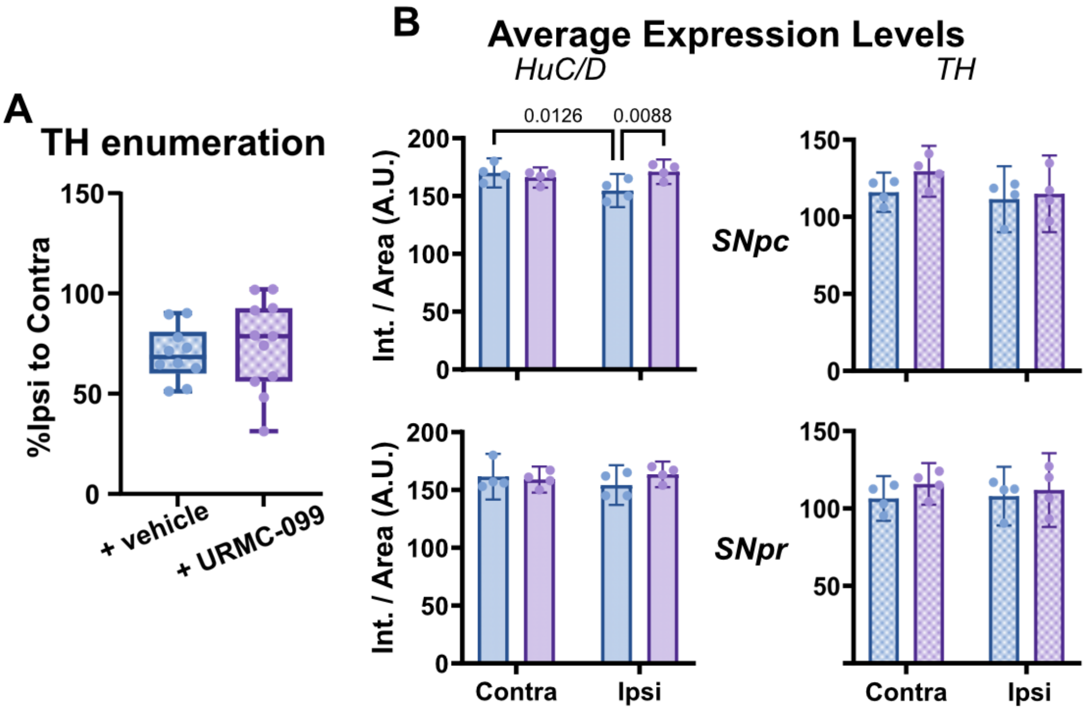

**<u>Figure S1. Extended data for Figures 1D and 1E relevant to neuronal integrity in the SNpc and SNpr.</u>**

**(A)** AI-based enumeration of TH+ cell bodies as a measure of total cell counts for dopaminergic neurons in the SN did not demonstrate differences between therapeutic cohorts. Both exhibit ~ 25% fewer counts of TH+ neurons in the ipsilateral hemisphere compared to the contralateral (n = 10).

**(B)** Quantification of average expression levels for all HuC/D+ and dopaminergic-specific TH+ neurons in the SNpc and SNpr. Statistical analyses were performed as full-effects two-way ANOVAs with Fisher's LSD post-hoc corrections (n = 4 for each group).

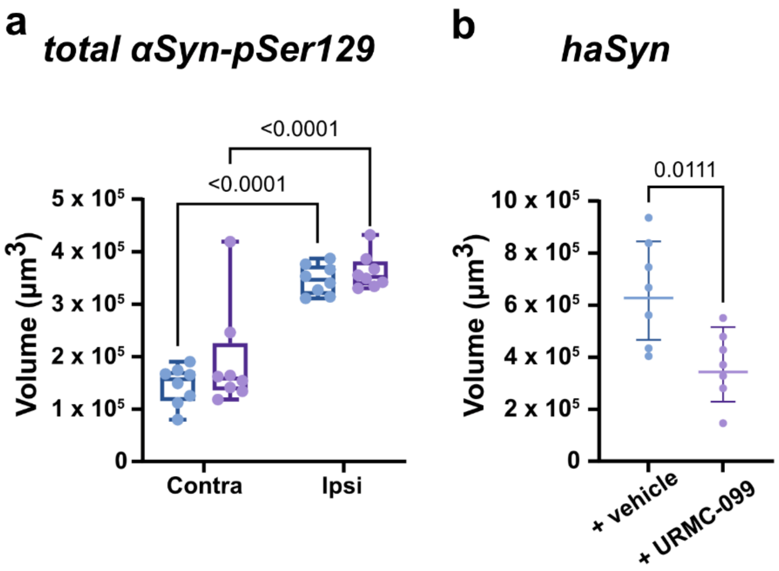

**Figure S2. Extended data for Figures 3A and 3B relevant to pathologic synuclein burden.**

Quantifications for (a) total αSyn-pSer129 burden and (b) total haSyn (Syn211+) burden by spatial distribution in SN (volume). Statistical analyses were performed as (a) a full-effects two-way ANOVA with Fisher's LSD post-hoc correction (n = 8 per group) and (b) an unpaired two-tailed t-test (n = 8 per group).

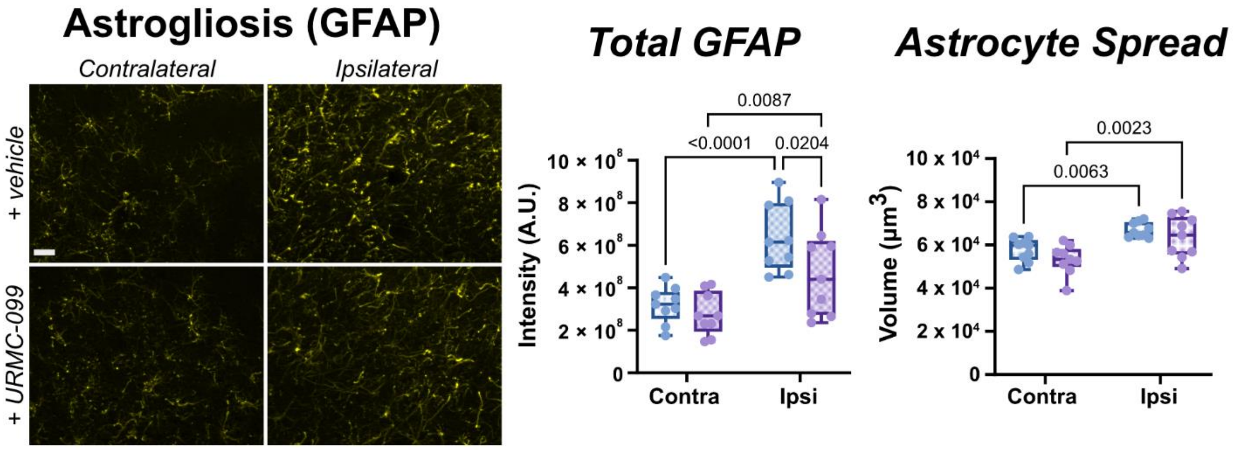

**Figure S3. Extended data for Figure 3D relevant to astrocytic activation in the SN.**

IF panels showing GFAP immunoreactivity in both contralateral and ipsilateral SN for rats given either vehicle or URMC-099 therapy. Scale bar is 30 µm. Quantifications for total GFAP expression (intensity) and astrocyte spread (volume). Statistical analyses were performed as full-effects two-way ANOVAs with Fisher's LSD post-hoc correction (n = 9 per group).

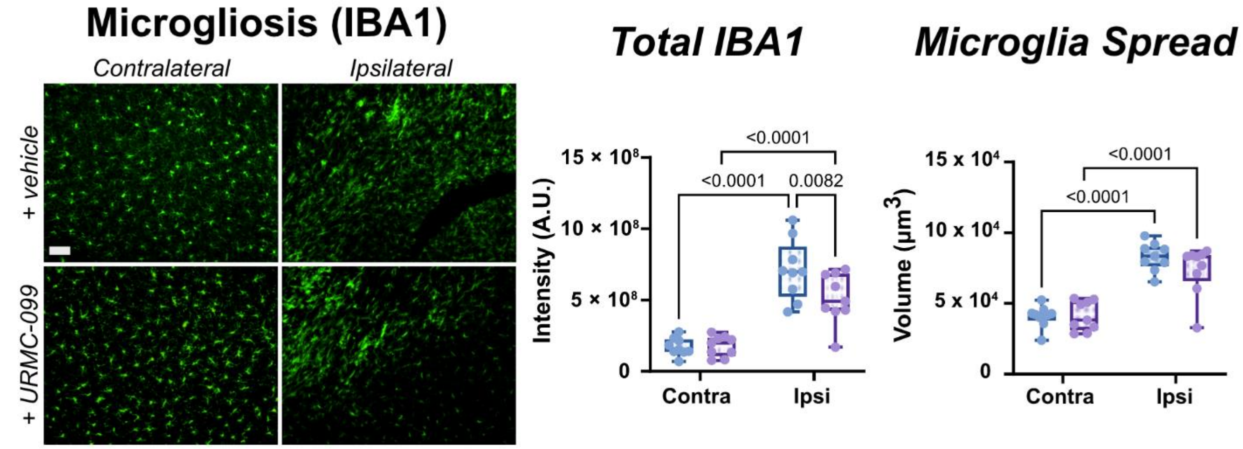

**<u>Figure S4. Extended data for Figure 3E relevant to microglia activation in the SN.</u>**

IF panels showing IBA1 immunoreactivity in both contralateral and ipsilateral SN for rats given either vehicle or URMC-099 therapy. Scale bar is 30 µm. Quantifications for total IBA1 expression (intensity) and microglia spread (volume). Statistical analyses were performed as full-effects two-way ANOVAs with Fisher's LSD post-hoc correction (n = 9 per group).

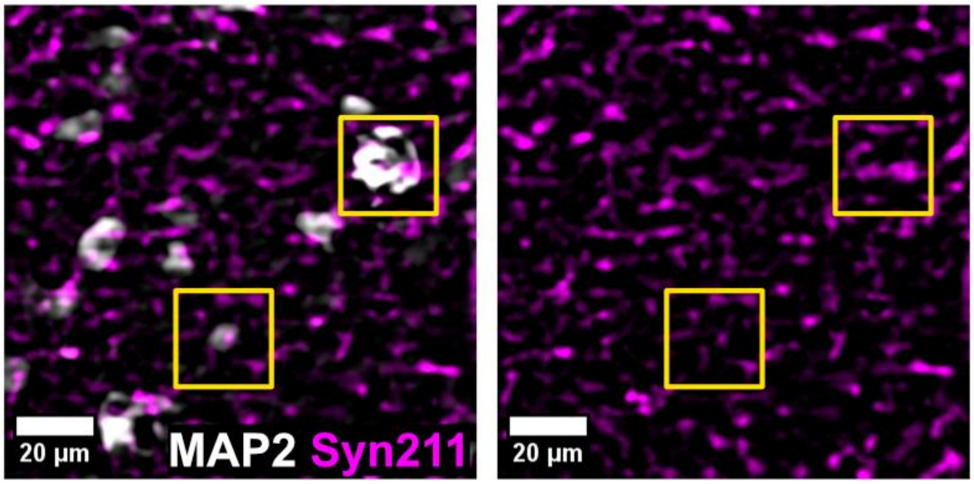

**<u>Figure S5. Somatic inclusions containing Syn211+ haSyn in MAP2+ neurons in the ipsilateral SN.</u>**

Bright MAP2+ soma from neurons in the SN were found to contain Syn211+ inclusions. Yellow boxes outline regions that demonstrate colocalization of Syn211+ features in MAP2+ neural bodies. Scale bar is 20 µm.

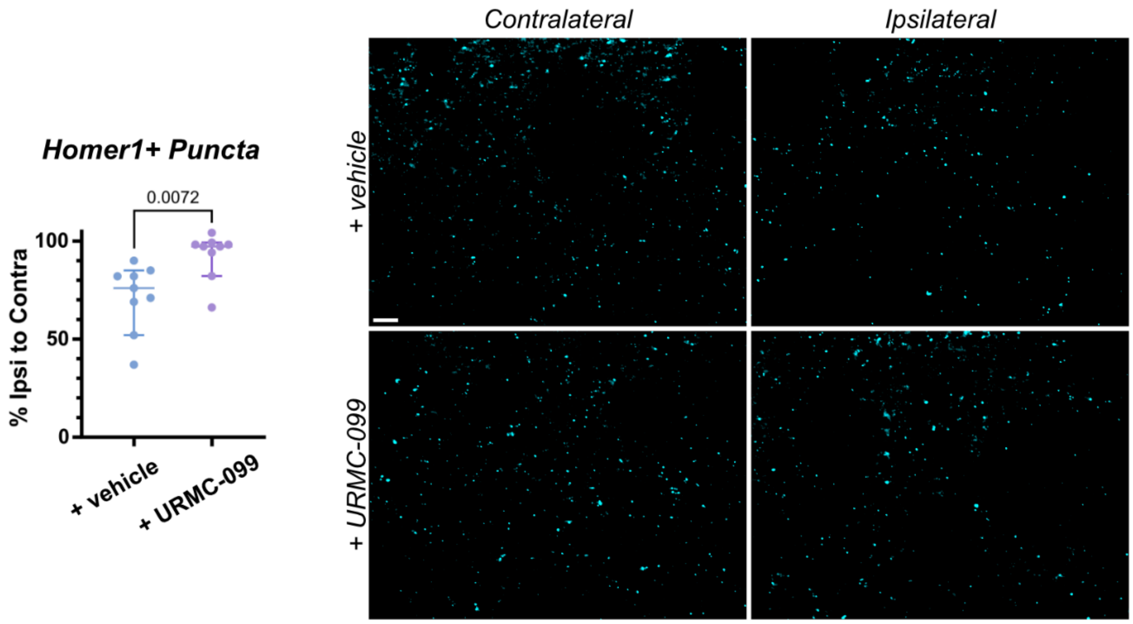

**Figure S6. Evaluation of neuroprotection in the striatum with URMC-099 treatment.**

IF panels for Homer1 immunoreactivity on synaptic densities in both contralateral and ipsilateral striatum for rats given either vehicle or URMC-099 therapy. Scale bar is 20 µm. Quantification for a neurodegenerative pathology was done by comparing the ratio of the average count of synaptic densities in the ipsilateral to the contralateral for each rat brain. Statistical analysis was performed as an unpaired two-tailed t-test (n = 8 per therapeutic group).

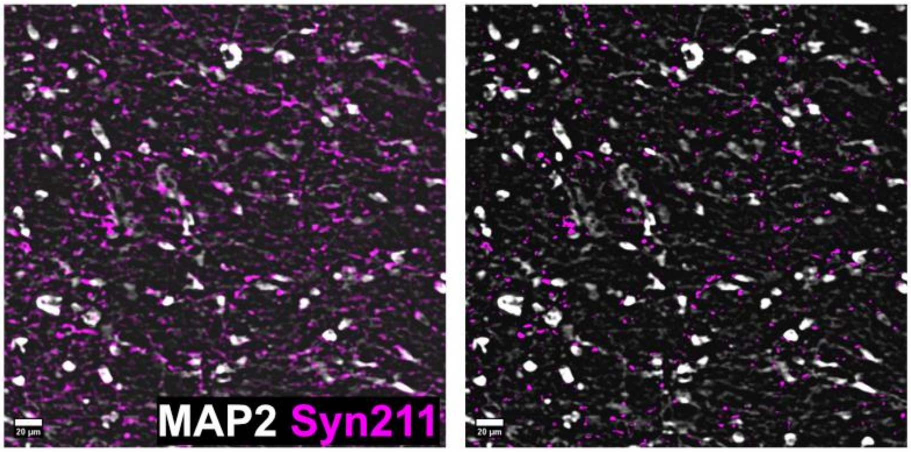

**Figure S7. Extended data for Figure 4F related to non-neuronal haSyn.**

IF panels representative of haSyn expression with respect to neurons in the ipsilateral SN. Left panel shows localization of all Syn211+ features with respect to all neurons labeled by MAP2. The intersection of the two channels was computed using binary logic functions for image calculation in ImageJ, followed by the XOR function of that intersection with the Syn211 channel to produce the set of Syn211+ features not contained in MAP2+ regions (right). This population represents non-neuronal detection of haSyn. Scale bars are 20 µm.

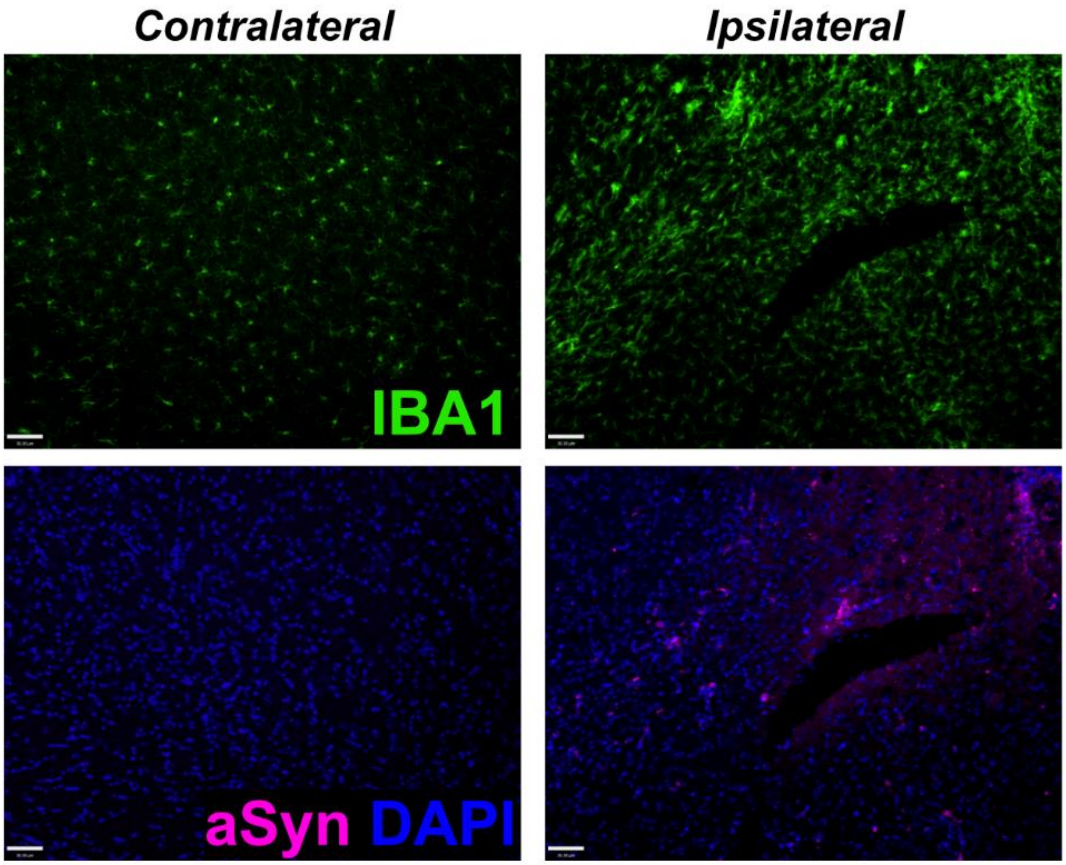

**Figure S8. Extended data for Figure 5D relevant to hemispheric localization of haSyn.**
IF panels representative of haSyn expression localized to the lesion-bearing ipsilateral SN with ensuant microglial activation via increased IBA1 reactivity. Scale bars are 30 μm.

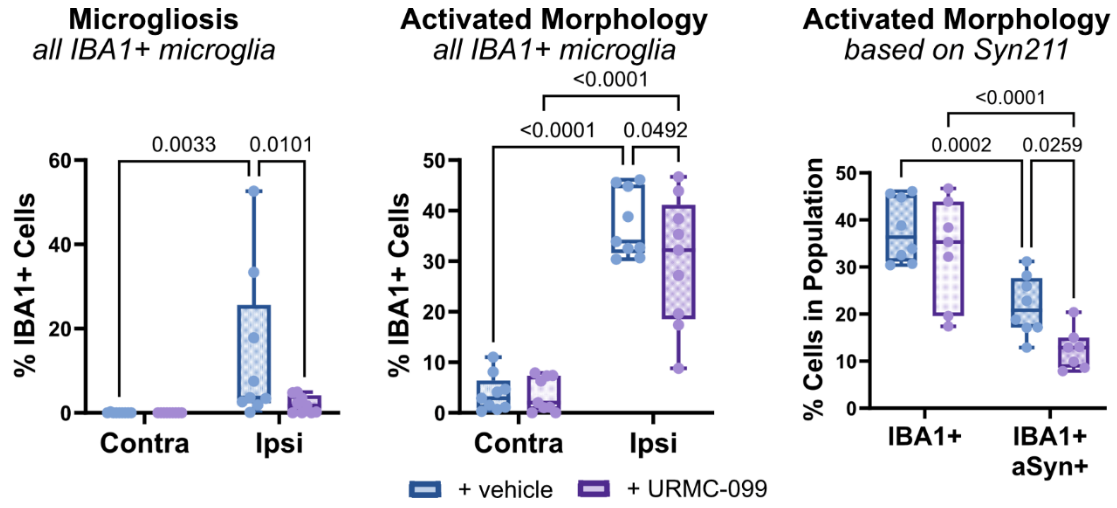

**Figure S9. Extended data for Figure 6 relevant to single-cell segmentation of IBA1+ microglia.**

Quantifications of nigral microglia based on population-based metrics obtained from single-cell segmentation and classification for activation state (IBA1 expression levels) and effector state (morphologies). **(Left)** Expanded quantification of microgliosis representing all IBA1+ cells in both hemispheres, demonstrating a significant ($p = 0.0033$) increase in the fraction of IBA1+ cells in a state of microgliosis in the ipsilateral SN that is significantly ($p = 0.0101$) attenuated with URMC-099 treatment. **(Middle)** Changes in overall “activated” morphology based on all IBA1+ cells across the SN in both hemispheres. This reflects the aggregate number of cells that were classified as either “hyper-ramified” or “amoeboid”. Both therapeutic groups demonstrate a significant increase ($p < 0.0001$) in the frequency of microglia demonstrating an activated morphology in the ipsilateral SN. URMC-099 reduces the net burden of morphologically active microglia ($p = 0.0492$). **(Right)** Activated morphology analysis based only on microglia in the ipsilateral SN that are divided based on dual positivity with Syn211+ haSyn. This demonstrates that upon engulfment of αSyn, IBA1+ Syn211+ cells demonstrate a decreased frequency in a non-homeostatic-like morphology ($p = 0.0002$ and $p < 0.0001$ for vehicle and URMC-099, respectively) compared to IBA1+ Syn211- cells. URMC-099 treatment demonstrates an additional reduction in the net fraction of morphologically active IBA1+ Syn211+ microglia ($p = 0.0259$). Statistical analyses were performed as full-effects two-way ANOVAs with Fisher’s LSD post-hoc correction ($n = 9$ per group).

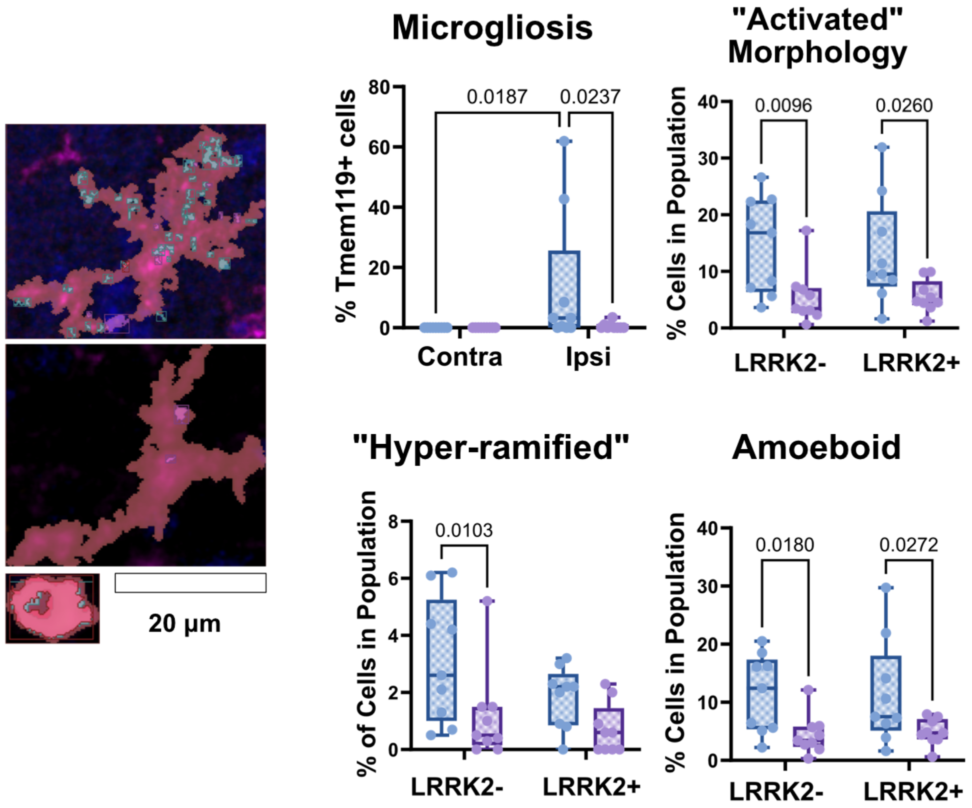

**Figure S10. Extended data for Figure 6 relevant to single-cell segmentation of Tmem119+ microglia.**

IF panels representative of segmented Tmem119+ microglia demonstrating different effector state morphologies. Quantifications of nigral Tmem119+ microglia using population-based metrics obtained from single-cell segmentation and classification. AAV-induced haSyn expression in the ipsilateral SN results in a population of microglia demonstrating microgliosis ($p = 0.0187$) that is attenuated by once-daily URMC-099 therapy ($p = 0.0237$). URMC-099 therapy also reduced the population of microglia that exhibited an activated, non-homeostatic-like effector state morphology regardless of LRRK2- ($p = 0.0096$) and LRRK2+ ($p = 0.026$) status. Comparison of Tmem119+ microglia based on heightened LRRK2 expression (i.e., LRRK2- or LRRK2+) did not demonstrate a clear dependency between LRRK2 activation levels and net presentation of effector state morphologies, indicating that simply inhibiting LRRK2 was not sufficient in maintaining dynamicity of the microglia. Statistical analyses were performed as full-effects two-way ANOVAs with Fisher's LSD post-hoc correction ($n = 9$ per group).

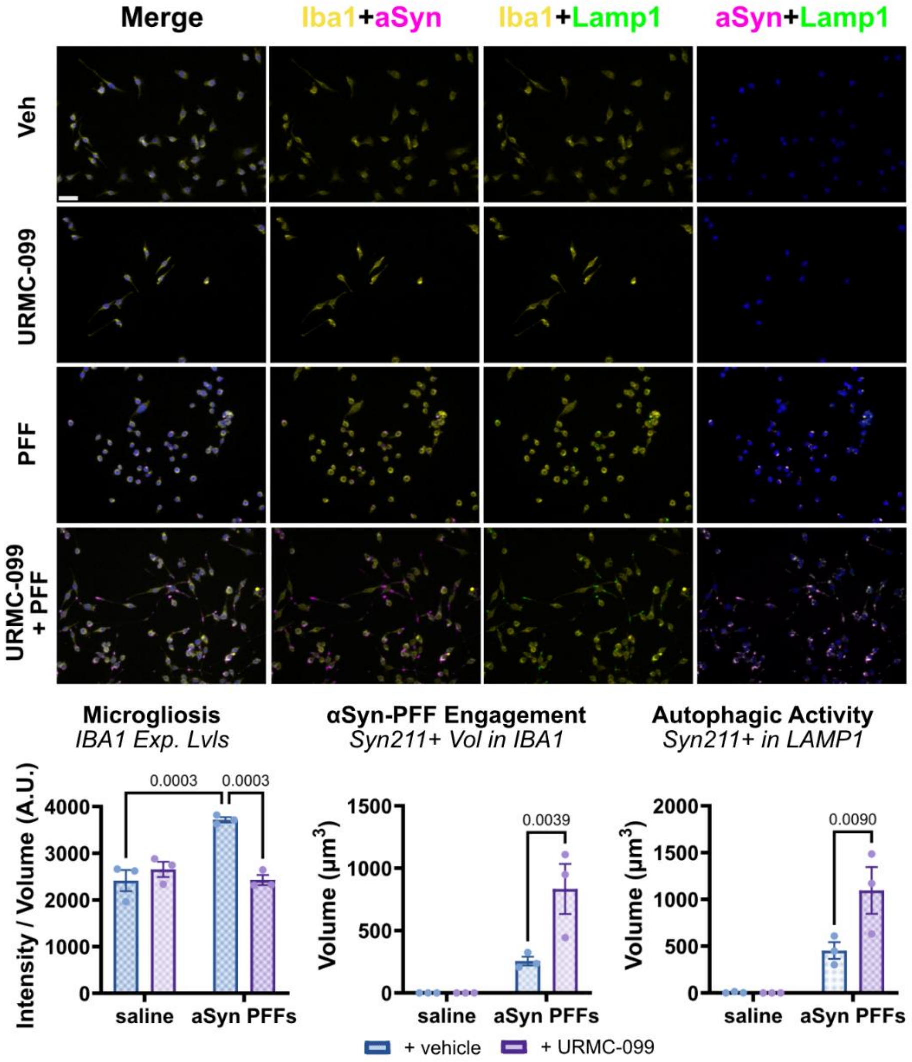

**Figure S11. Extended data for Figure 7 relevant to in vitro BV-2 model for αSyn PFF related stress.**

**(Top)** ICC panels of IBA1+ microglia examining changes in morphology, IBA1 expression levels, and autophagolysosomal activity based in CD68 expression and engulfment of Syn211+ PFFs. Scale bar is 20 μm. Treatment groups were: veh. (0.1% v/v saline + 0.1% v/v DMSO); URMC-099 (0.1% v/v saline + 100 nM URMC-099); PFF (2 μg/mL αSyn PFF + 0.1% v/v DMSO); and URMC-099 + PFF (2 μg/mL αSyn PFF + 100 nM URMC-099). **(Bottom)** Quantifications for microglioisis, level of αSyn-PFF burden, and autophagic activity. Exposure to PFFs significantly ($p = 0.0003$) induced microgliosis compared to saline-matched vehicle control. This was mitigated by co-treatment with URMC-099 ($p = 0.0003$) to maintain average IBA1 expression levels similar to saline-matched negative controls. No false positives for Syn211+ PFF signal were detected in the analysis pipeline for saline-matched negative controls. URMC-099 treatment doubled the average amount of PFFs the microglia were in contact with ($p = 0.0039$) with a similar increase in the engulfment of PFFs in CD68+ structures ($p = 0.0090$). Statistical Analyses were performed as two-way ANOVAs with Fisher's LSD post-hoc correction ($n = 3$ per group).

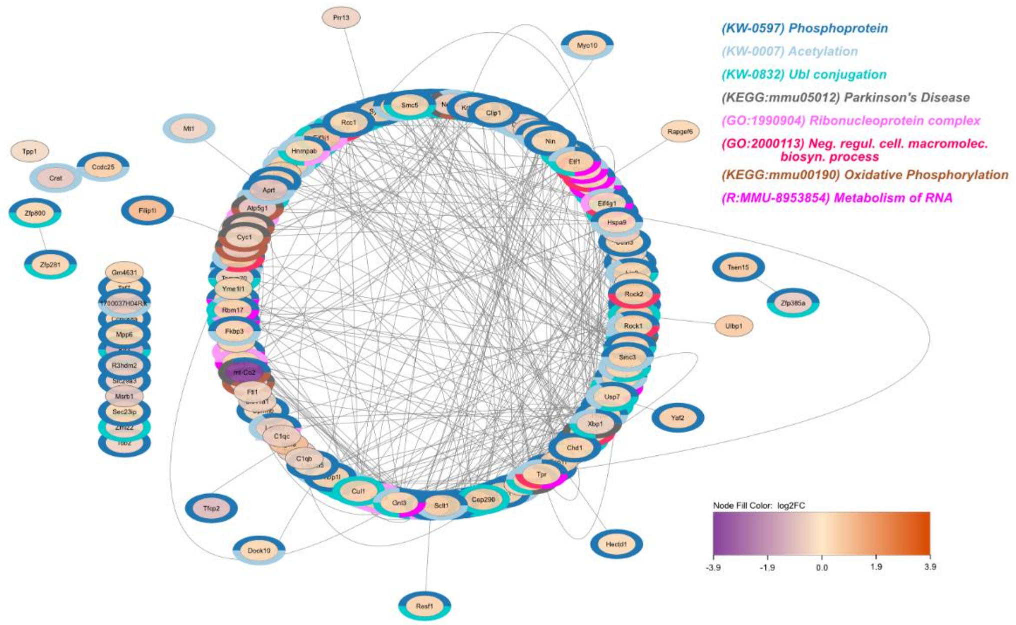

**Figure S12. Extended data for Figure 7D relevant to transcriptomic analysis of αSyn-PFF + DMSO vehicle treated BV-2 compared to saline + DMSO negative control.**

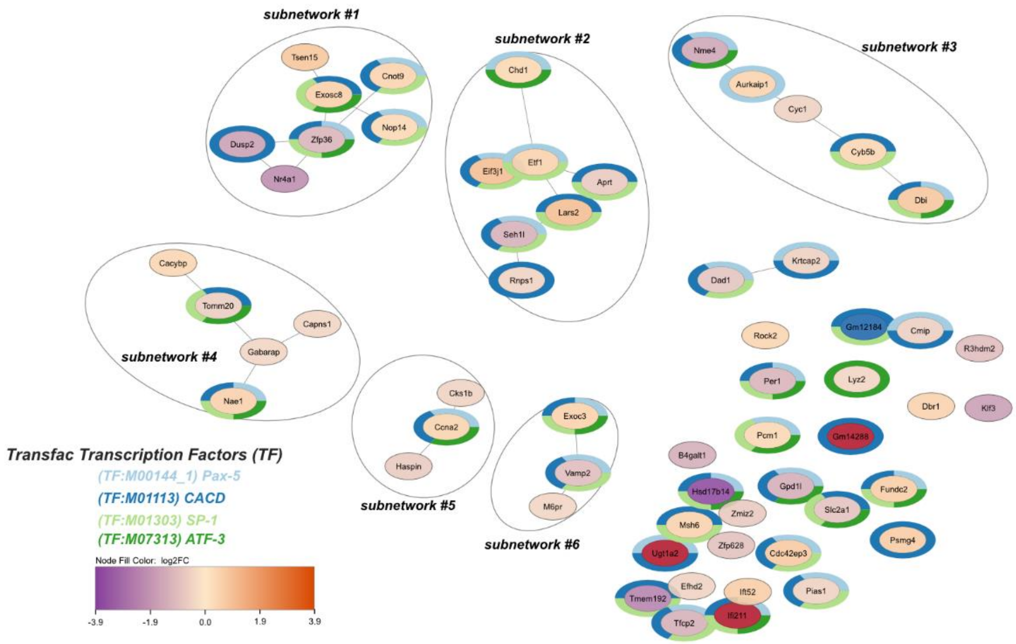

**Figure S13. Extended data for Figure 7D relevant to transcriptomic analysis of αSyn-PFF + URMC-099 treated BV-2 compared to full vehicle (saline + DMSO) control.**

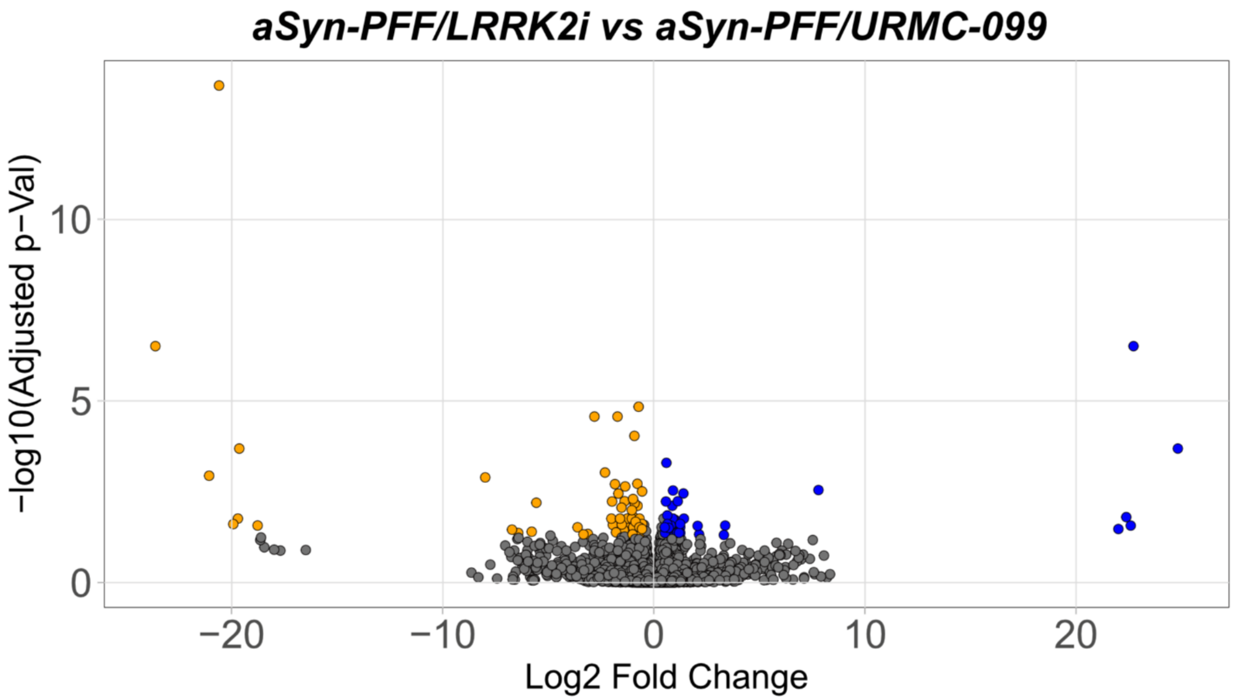

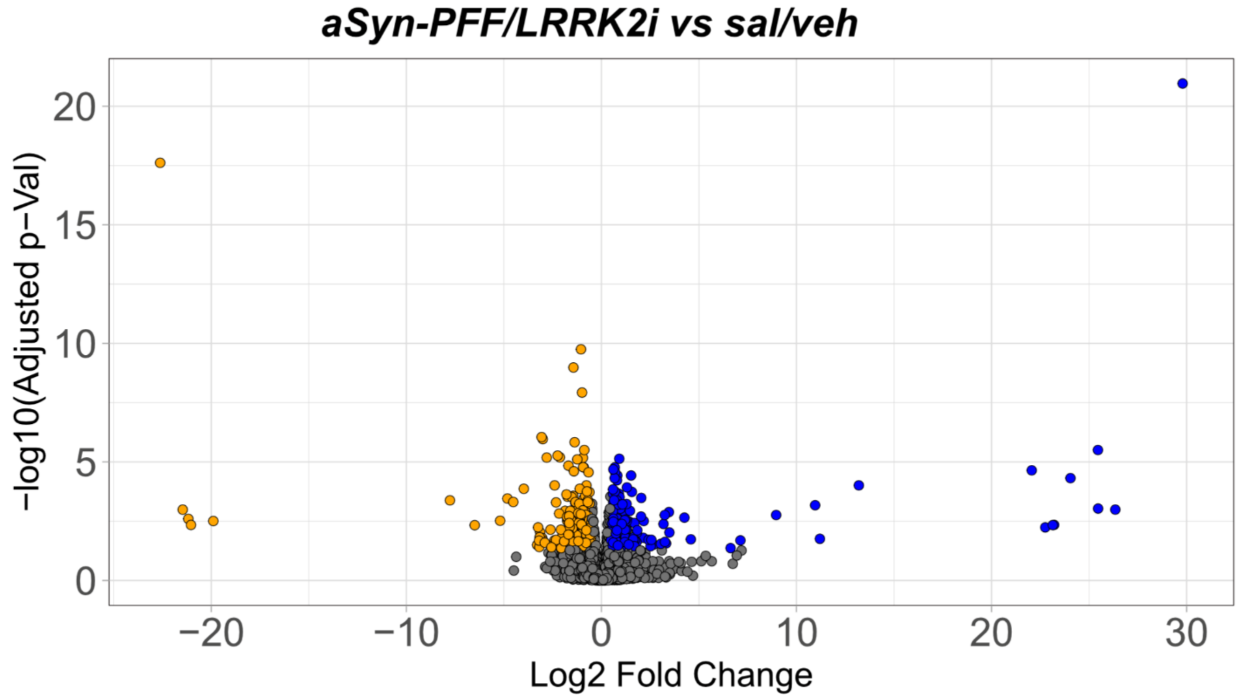

**Figure S14. Extended data for Figure 7F relevant to differences between solo inhibition of LRRK2 compared to dual MLK3/LRRK2 inhibition by URMC-099 in αSyn-PFF treated BV-2.**

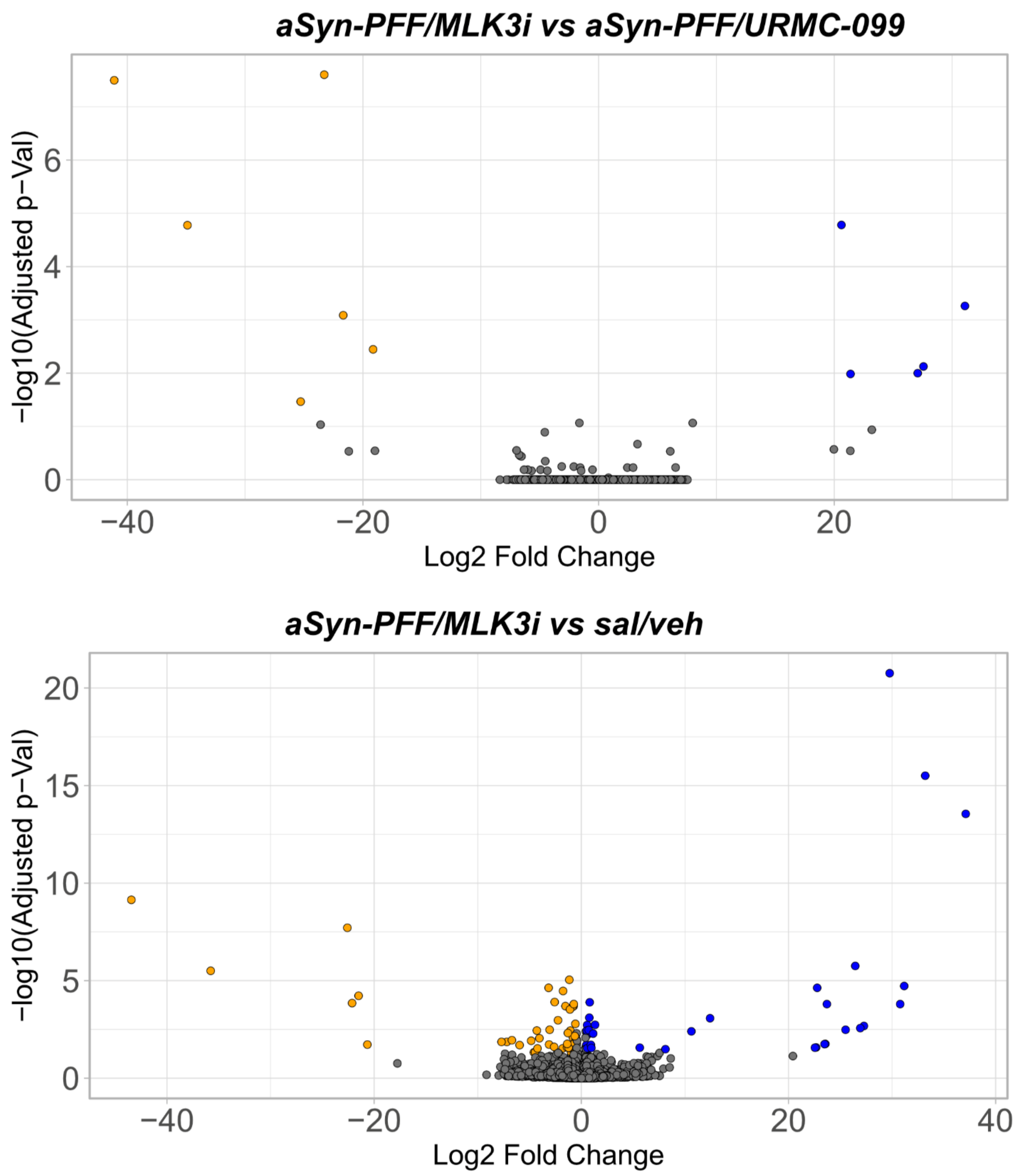

**Figure S15. Extended data for Figure 7F relevant to differences between solo inhibition of MLK3 compared to dual MLK3/LRRK2 inhibition by URMC-099 in αSyn-PFF treated BV-2.**

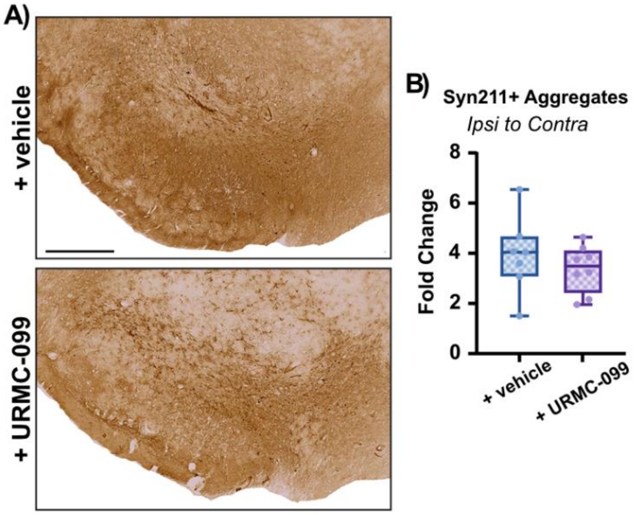

**Figure S16. Proteinase K resistant SYN211+ deposits in the substantia nigra**

**(A)** Brightfield images of DAB-stained SYN211 immunoreactive aggregates in the substantia nigra following proteinase K treatment. **(B)** Quantification of fold-changes in the presence of Syn211+ aggregates in the ipsilateral hemisphere compared to the contralateral. No differences were observed between therapeutic cohorts. Statistical Analysis was performed as a two-tailed non-parametric Mann-Whitney test.

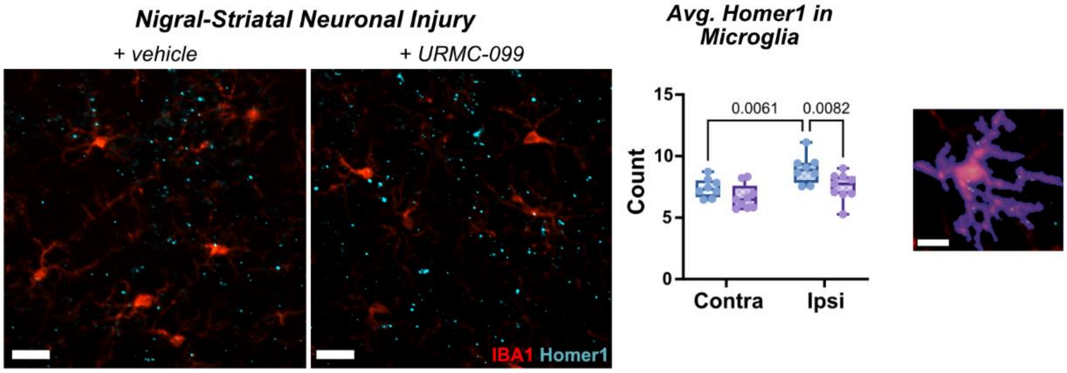

**Figure S17. Extended data for Figure S6 related to microglia pruning of Homer1+ synapses in striatum.**

**(Left)** IF panels for IBA1+ microglia contacting Homer1+ synapses in ipsilateral striatum for rats given either vehicle or URMC-099 therapy. Scale bar is 20 μm. **(Middle)** Following single-cell segmentation of IBA1+ cells, the microglia in the lesion-bearing ipsilateral hemisphere on average demonstrated a ~30% increase in the number of synapses they were contacting at a per cell basis ($p = 0.0061$). This was prevented by URMC-099 therapy ($p = 0.0082$). **(Right)** Example of segmented IBA1+ microglia with contacted Homer1+ synapses (yellow stars). Scale bar is 5 μm. Statistical analysis was performed as a two-way ANOVA with Fisher's post-hoc correction ($n = 9$).
